# Mechanosensing and IL-13 Signaling Synergistically Modulate Intestinal Stem Cell Differentiation via STAT6 and YAP

**DOI:** 10.64898/2026.02.19.706676

**Authors:** Sarbari Saha, Thao Nguyen, Cornelis Mense, Marie Touzet-Robin, Karen Kresbach, Stephan A. Eisler, Ulrich S. Schwarz, Andrew G. Clark

## Abstract

A long-term complication of chronic inflammation is the mechanical stiffening of the tissue, culminating in fibrosis. Fibrosis can severely disrupt tissue function and is a major risk factor for other diseases. It is not currently well understood how fibrosis impacts the response to inflammatory signals. To address this, we investigated cross-talk between cellular mechanosensing and the response to Interleukin (IL)-13, a cytokine associated with inflammatory bowel diseases (IBDs). Using 3D intestinal organoids and organoid monolayer culture, we uncovered a synergy between mechanosensing and IL-13 signaling in regulating intestinal stem cell differentiation. Through quantitative high-resolution microscopy and functional inhibition, we found that this response requires activation of STAT6, a known mediator of IL-13. Both IL-13 and high substrate stiffness increase cellular traction forces and focal adhesion formation, but at the expense of reduced tension at cell-cell junctions and compromised epithelial barrier function. The mechanosensing and IL-13 responses require actomyosin contractility and YAP, which is activated downstream of a positive feedback loop involving STAT6-dependent myosin-2 activation. Our results establish a novel STAT6-YAP signaling axis that integrates inflammatory and mechanical cues to regulate intestinal cell fate and barrier integrity, opening new avenues to target epithelial dysfunction in fibrosis, chronic inflammation and regenerative medicine.

## Introduction

The intestinal epithelium undergoes continual self-renewal due to intestinal stem cell proliferation. Intestinal stem cells differentiate into specialized cell lineages, primarily absorptive enterocytes, but also various secretory cell types including goblet cells, enteroendocrine cells, and tuft cells. Cell fate specification is a crucial mechanism for maintaining epithelial homeostasis, enabling the tissue to support regeneration and respond to environmental stressors^1,2^. Differentiation is governed by an intricate interplay between intrinsic transcriptional programs and extrinsic environmental inputs, including cytokine signaling and mechanical cues from the surrounding extracellular matrix (ECM)^3–5^.

During chronic inflammation, for example in inflammatory bowel diseases (IBDs), normal cell type specification patterns become disrupted. This can result in the depletion of goblet cells in IBDs and expansion of secretory cells in colorectal cancer^6,7^. IBDs are associated with increased levels of inflammatory cytokines (ICs) including Tumor Necrosis Factor (TNF)-α, Interleukin (IL)-6, IL-13, and Interferon (IFN)-γ. IC signaling can interfere with both transcriptional and ECM-mediated signals, ultimately impairing intestinal stem cell renewal and barrier function^8,9^. This promotes a cycle of tissue damage, dysregulated repair and disease progression^10–12^. Sustained exposure to ICs also stimulates stromal and immune cells in the underlying tissue to increase secretion and remodeling of local ECM networks, ultimately leading to fibrosis^13,14^. Understanding how immune and mechanical signals interact to regulate epithelial homeostasis is thus critical for understanding IBD pathogenesis and uncovering novel therapeutic targets.

Among IBD-associated ICs, IL-13 plays a pivotal role in epithelial reprogramming^15^. IL-13 is elevated in the intestine in conditions such as ulcerative colitis and parasitic infections and also in other tissues such as asthmatic lung^16–18^. Upregulation of IL-13 signaling promotes goblet cell hyperplasia, increases mucus secretion and disrupts barrier integrity by dysregulating tight junctions^19^. IL-13 signaling is mediated primarily through the activation of Signal Transducer and Activator of Transcription 6 (STAT6)^18,20,21^. IL-13 binds predominantly to transmembrane IL-13Rα1-IL-4Rα heterodimers, leading to phosphorylation of STAT6-Y641 by Janus Kinase (JAK) proteins. According to the canonical IL-13 signaling pathway, phosphorylated STAT6 (pSTAT6) proteins dimerize and accumulate in the nucleus where they act as transcription factors, promoting expression of cytokines and genes involved in cell cycle progression and ECM protein production^22^. How IL-13 signaling is modulated by the mechanical properties of the tissue microenvironment remains less understood.

Recent findings suggest that increased microenvironment stiffness, characteristic of IBD-linked fibrosis, shifts intestinal epithelial stem cell fate. Increased substrate stiffness diminishes the LGR5⁺ stem cell pool and promotes goblet cell differentiation^8^. Stiffness-dependent modulation of IL-13/STAT6 signaling has been observed in immune cells. In macrophages, substrate rigidity enhances STAT6 nuclear translocation, promoting M2 polarization in response to IL-13^23^. Complementing this, the mechanosensitive ion channel Piezo1 modulates IL-13-induced STAT6 activation on stiff substrates by stimulating Ca^2+^ influx, implicating biomechanical signaling in cytokine responsiveness^24^. These findings suggest that IL-13 signaling and mechanosensing could behave in a synergistic or co-dependent manner. However, this behavior has been described in individual immune cells, and the potential synergies between mechanosensing and cytokine signaling in polarized epithelia are currently not understood.

Mechanical stimuli have a profound impact on actomyosin contractility, which plays a central role in epithelial morphogenesis, remodeling, and barrier function^25^. IL-13 is known to influence actin cytoskeletal organization in epithelial cells, particularly in pathological conditions such as asthma and hepatic fibrosis, where IL-13 drives goblet cell metaplasia and tissue stiffening through myofibroblast activation and enhanced actomyosin contractility^15,26,27^. These effects are mediated in part by RhoA signaling and myosin light chain (MLC) phosphorylation, underscoring the importance of IL-13 in cytoskeletal regulation^28^. However, the molecular mechanisms mediating crosstalk between IL-13/STAT6 signaling and actomyosin remodeling in epithelial cells is currently not well defined.

Mechanical cues from the microenvironment are integrated at cell-cell and cell-substrate adhesions, where they regulate cytoskeletal organization, barrier integrity, and epithelial morphology^29,30^. Intestinal epithelial cells generate compartmentalized traction forces that are transmitted via focal adhesion (FA) proteins such as vinculin and paxillin as well as through junctional complexes including adherens junctions (AJs), tight junctions (TJs), and desmosomes^31–34^. These adhesion structures coordinate actomyosin-generated forces to regulate tissue architecture, collective migration, and cell fate decisions through regulation of mechanosensory transcriptional programs^35,36^. One of the most prominent mechanosensory transcription factors is Yes-associated protein (YAP) and its paralog Transcriptional co-activator with PDZ-binding motif (TAZ), which regulate various downstream processes including epithelial differentiation^37^. In colorectal cancer cells and intestinal organoids, high substrate stiffness induces YAP nuclear translocation, leading to matrix remodeling, stem cell depletion, and goblet cell differentiation^8,38^. Moreover, YAP activity is modulated by intestinal stem cell niche factors such as Wingless-related integration site (Wnt), Bone Morphogenetic Protein (BMP) and lipolysis-stimulated lipoprotein receptor (LSR), which influences secretory cell differentiation^39^. In addition to YAP, mechanosensitive Piezo channels have recently been shown to regulate intestinal stem cell renewal and differentiation during homeostasis^40^.

Taken together, evidence from previous studies suggest a highly integrated network wherein microenvironment stiffness, actomyosin contractility, cell adhesions and cytokine signaling converge to direct epithelial cell fate, function, and pathology. In this study, we address the crosstalk between mechanosensing and inflammatory signaling directly by investigating how substrate stiffness modulates the intestinal epithelial response to IL-13. Our data suggest that intermediate substrate stiffness optimally supports IL-13-induced goblet cell hyperplasia via STAT6 activation, while high substrate stiffness alone can mimic inflammatory signaling by promoting STAT6 nuclear localization and secretory differentiation even in the absence of inflammatory signaling. Actomyosin contractility links matrix stiffness and IL-13 signaling to STAT6 and YAP activation, both of which are essential for secretory cell fate decisions. Functional inhibition experiments suggest that STAT6 activation is upstream of YAP activation. To explain this, we propose a mechanism by which STAT6 drives expression of myosin light chain kinase 1 (MLCK1), which in turn phosphorylates MLC, increasing actomyosin contractility to promote YAP-mediated mechanotransduction. Together, our findings identify a STAT6- and YAP-dependent signaling axis that integrates inflammatory and mechanical signals to regulate intestinal epithelial differentiation, with implications for understanding tissue remodeling in fibrotic and inflammatory diseases.

## Results

### Substrate Stiffness Sensitizes the Intestinal Epithelium to IL-13 and Promotes Goblet Cell Hyperplasia via STAT6

While IL-13 is a known inducer of goblet cell differentiation, recent evidence suggests that increased substrate stiffness can also drive changes in cell fate specification. How mechanical cues and inflammatory signals potentially interact to regulate epithelial cell fate remains poorly understood. To address this, we used mouse intestinal organoids, which recapitulate various aspects of intestinal homeostasis *in vitro*, including epithelial stem cell proliferation and differentiation^41,42^. To investigate the role of microenvironment stiffness, we cultured mouse intestinal organoid monolayers on polyacrylamide (PAA) gels with varying stiffness levels: soft (0.5 kPa), intermediate (5 kPa) and stiff (15 kPa). These stiffness levels were chosen to mimic the mechanical properties of healthy and fibrotic tissue environments^14^. In addition, previous studies have indicated that 5kPa represents a critical stiffness threshold for mechanosensing^43^. Organoid monolayers were then treated with mouse recombinant IL-13 and stained with the fluorescently-tagged lectin Ulex europaeus agglutinin I (UEA I), which labels intestinal mucus and goblet cells. We then visualized monolayers using confocal microscopy and quantified goblet cell frequency using a customized image analysis pipeline (Supplementary Figure 1). On soft substrates, IL-13 treatment had no effect on the fraction of goblet cells (Fig. 1A, B). For monolayers on intermediate stiffness PAA gels, IL-13 significantly increased goblet cell frequencies from approximately 15% to 40%, while on stiff substrates, the frequency of goblet cells was elevated even in the absence of IL-13 treatment, and additional IL-13 treatment did not further change goblet cell frequency. Together, these data suggest that monolayers on intermediate stiffness substrates are mechanically primed to respond to IL-13, while high substrate stiffness alone mimics an IL-13-like response.

**Figure 1.**
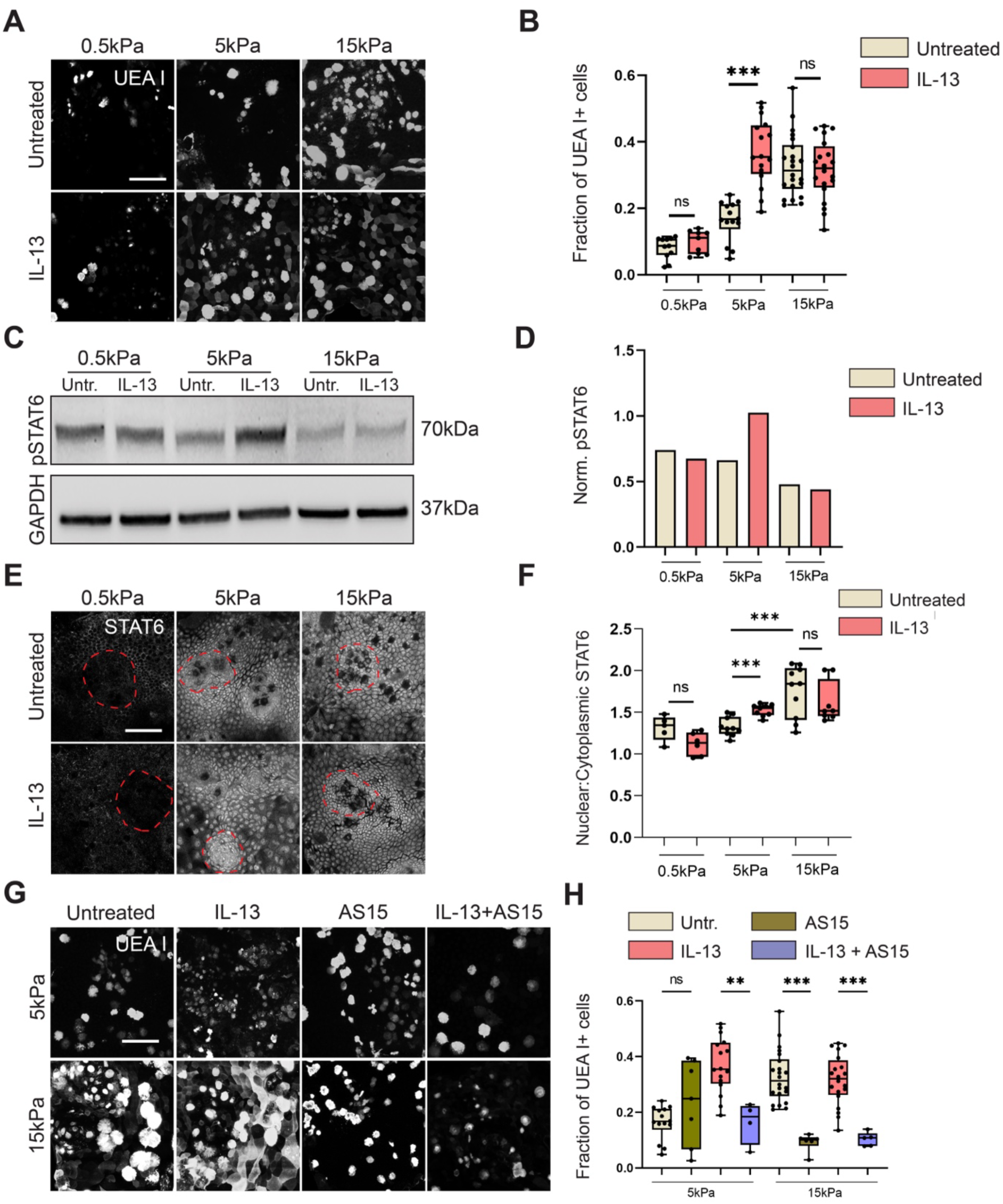
Substrate Stiffness Sensitizes the Intestinal Epithelium to IL-13 and Promotes Goblet Cell Hyperplasia via STAT6. **A.** Representative micrographs of organoid monolayers cultured on soft (0.5 kPa), intermediate (5 kPa), or stiff (15 kPa) polyacrylamide (PAA) gels, treated with or without IL-13 and stained with fluorescently tagged UEA I to identify goblet cells. Images are max projections of z-stacks. SB: 20µm. **B.** Quantification of the fraction of UEA I+ cells for conditions as in *A*. Each dot represents one organoid monolayer from *n* = 11, 9, 14, 16, 22, 20 organoid monolayers and *N* = 3, 3, 3, 3, 3, 3 independent experiments. **C.** Western blot of phosphorylated STAT6 (pSTAT6) from organoid lysates after culture on substrates of varying stiffness with or without IL-13 treatment. **D.** Densitometric quantification of pStat6 levels (normalized to GAPDH) from Western blot in *C*. Bar plot represents mean of triplicate quantification. **E.** Representative micrographs of organoid monolayers cultured on soft/intermediate/stiff PAA gels, treated with or without IL-13 and immunostained for STAT6. Red dashed lines indicate the crypt-like compartments. SB: 20µm. **F.** Quantification of nuclear to cytoplasmic STAT6 fluorescence intensity ratio for conditions in *E*. Each dot represents one organoid monolayer from *n* = 5, 6, 10, 9, 10, 8 organoid monolayers and *N* = 3, 3, 3, 3, 3, 3 independent experiments. **G.** Representative micrographs images of organoid monolayers cultured on intermediate/stiff PAA gels, treated with IL-13 and/or the STAT6 inhibitor AS1517499 (AS15) and stained with fluorescently tagged UEA I to identify secretory goblet cells. Images are max projections of z-stacks. SB: 20µm. **H.** Quantification of the fraction of UEA I+ secretory cells under different stiffness and treatment conditions as in *G*. Each dot represents one organoid monolayer from *n* = 14, 7, 16, 4, 22, 6, 20, 5 organoid monolayers and *N* = 3, 3, 3, 3, 3, 3, 3, 3 independent experiments. The untreated and IL-13 data are pooled from the same data plotted in *1B*. For panels B, F, H, One-way ANOVA was performed (p = 8.3e-19, 1.2e-6, 2.7e-12). Pairwise tests performed using Tukey’s HSD post-hoc text (**p<0.01, ***p<0.001, ns:p>0.05).

These data prompted us to investigate the broader impact of IL-13 on differentiation during epithelial homeostasis. To assess this, we treated 3D organoids with IL-13 and quantified the number of crypt-like buds in individual organoids. Organoids treated with IL-13 had a significantly higher number of buds compared to untreated organoids (Supplementary Figure 2A,B). As the crypt forming ability was compromised in IL-13 treated organoids, we next sought to investigate the proliferative potential of intestinal organoids upon treatment with IL-13 by immunostaining for Ki67. In untreated control organoids, Ki67 expression was strictly localized to the buds, consistent with the established compartmentalization of proliferative cells within these crypt-like structures^41,42^. IL-13 treatment led to a significant expansion of Ki67+ cells with aberrant spatial distribution throughout the entire organoid structure (Supplementary Figure 2C,D). To determine how the organization of the intestinal stem cell compartment is influenced by substrate mechanics, we cultured organoid monolayers on intermediate (5 kPa) and stiff (15 kPa) substrates, treated with IL-13 and immunostained for Olfm4 to label intestinal stem cells. On intermediate stiffness, IL-13 treatment led to a reduction in the stem cell compartment area (Supplementary Figure 2E,F). On stiff substrates, the area of the stem cell compartment was already reduced even in the absence of IL-13; additional IL-13 treatment on stiff substrates did not further influence stem cell compartment area. Together, these results suggest that both increased substrate stiffness and IL-13 signaling not only drive secretory cell differentiation but also restrict the stem cell niche.

The interdependence of IL-13 and substrate stiffness suggests a synergy between mechanosensing and IL-13 signaling pathways. We next assessed how these cues interact to control JAK-STAT6 activation by investigating STAT6 phosphorylation and nuclear translocation. Interestingly, previous studies indicate that phosphorylation may not be required for STAT6 nuclear translocation in certain cell types^44,45^. To investigate this, we cultured organoids on polyacrylamide (PAA) gels with varying stiffness (soft, intermediate, and stiff) with or without IL-13 treatment. Organoid lysates were subsequently harvested, and pSTAT6 levels were measured via Western blot (Fig. 1C). Baseline pSTAT6 levels decreased with increasing substrate stiffness in untreated organoid monolayers. In line with the goblet cell differentiation data, IL-13 treatment induced a significant increase in pSTAT6 levels only in organoids cultured on intermediate stiffness substrates, again suggesting a mechanical priming on intermediate stiffness (Fig. 1C,D). In parallel, we performed confocal microscopy and quantitative image analysis to assess the nuclear-to-cytoplasmic localization of total STAT6. Monolayers cultured on soft substrates did not display significant STAT6 nuclear localization with or without IL-13 treatment, whereas IL-13 treatment induced nuclear accumulation of STAT6 on intermediate stiffness substrates. On stiff substrates, the nuclear:cytoplasmic ratio of STAT6 increased even in the absence of IL-13 (Fig. 1E,F), suggesting that substrate stiffness alone can drive STAT6 nuclear translocation. To verify the localization of STAT6 in 3D organoids, we treated with IL-13 and immunostained for STAT6. Untreated control organoids displayed diffuse localization of STAT6. Upon treatment with IL-13, the nuclear:cytoplasmic ratio of STAT6 increased (Supplementary Figure 2G,H). These data suggest a mechanistic interplay between IL-13 signaling and mechanotransduction pathways, where intermediate stiffness substrates mechanically prime organoids for STAT6 activation, while on stiff substrates, nuclear STAT6 accumulation occurs independently of phosphorylation, indicating that mechanical cues can regulate STAT6 activation in the absence of cytokine signaling.

To determine whether STAT6 activation is required for mechanosensitive- and IL-13-mediated changes in cell fate specification, we next investigated whether inhibition of STAT6 could suppress goblet cell hyperplasia induced by either IL-13 or substrate stiffness. To this end, we treated organoid monolayers cultured on intermediate stiffness substrates (5 kPa) with IL-13 in the presence or absence of AS1517499 (AS15), a small molecule inhibitor that blocks STAT6 phosphorylation and downstream IL-4/IL-13 signaling^46^. While AS15 has been shown to inhibit STAT6 phosphorylation, its ability to interfere with potentially phosphorylation-independent functions of STAT6 has not been previously examined. AS15 treatment significantly reduced goblet cell frequency in IL-13-treated monolayers on intermediate stiffness substrates, effectively rescuing IL-13-induced goblet cell hyperplasia (Fig. 1G,H). To determine whether STAT6 inhibition also mitigates stiffness-induced goblet cell hyperplasia, which our previous data suggest occurs independently of STAT6 phosphorylation (Fig. 1C), we treated organoids on stiff substrates (15kPa) with AS15 in the absence of IL-13. Indeed, AS15 treatment significantly reduced mechanosensing- and IL-13 induced goblet cell frequency in organoid monolayers (Fig. 1G,H), indicating that STAT6 activity contributes to goblet cell differentiation downstream of mechanical cues, potentially through phosphorylation-independent mechanisms. Together, these findings suggest that both substrate stiffness and IL-13 promote goblet cell differentiation via STAT6-dependent pathways. Pharmacological inhibition of STAT6 is sufficient to suppress goblet cell hyperplasia driven by either inflammatory or mechanical stimuli, highlighting STAT6 as a central integrator of cytokine signaling and mechanotransduction in the regulation of epithelial cell fate.

### Actomyosin Contractility Links Substrate Stiffness and IL-13 to STAT6 Activation in Secretory Cells

Having established that IL-13 and substrate stiffness synergistically drive secretory cell differentiation via STAT6, we next sought to investigate how mechanical signals are transduced to influence STAT6 activity and epithelial cell fate. Given the critical role of the cytoskeleton in sensing and responding to substrate mechanics, we hypothesized that actomyosin contractility may be involved in linking substrate stiffness with STAT6 activation. To explore this, we investigated phosphorylated myosin light chain (pMLC) localization in organoid monolayers on substrates with different stiffness with or without IL-13. pMLC is a well-established key regulator of actomyosin contractility and serves as a downstream effector of mechanosensing pathways, driving changes in cytoskeletal tension and cell shape^47^. Following immunostaining for pMLC, we quantified relative cortical enrichment of pMLC by linescan analysis (Supplementary Figure 3A). In organoids cultured on intermediate stiffness PAA gels, IL-13 treatment increased pMLC enrichment at the cell cortex and the basal membrane of secretory cells (Figure 2A,B). On stiff substrates, pMLC exhibited strong cortical and basal localization in secretory cells even in the absence of IL-13. Addition of IL-13 on stiff substrates had no further effect on pMLC localization. In contrast to pMLC, organoid monolayers on intermediate stiffness or stiff substrates with or without IL-13 treatment displayed no significant changes in cortical or basal enrichment of Non-Muscle Myosin Heavy Chain 2A (MYH9) in secretory cells (Supplementary Figure 3B,C). These data indicate that both mechanical cues and IL-13 regulate pMLC, but not MYH9, localization, mirroring our earlier findings in which IL-13 sensitivity was optimal on intermediate stiffness.

**Figure 2.**
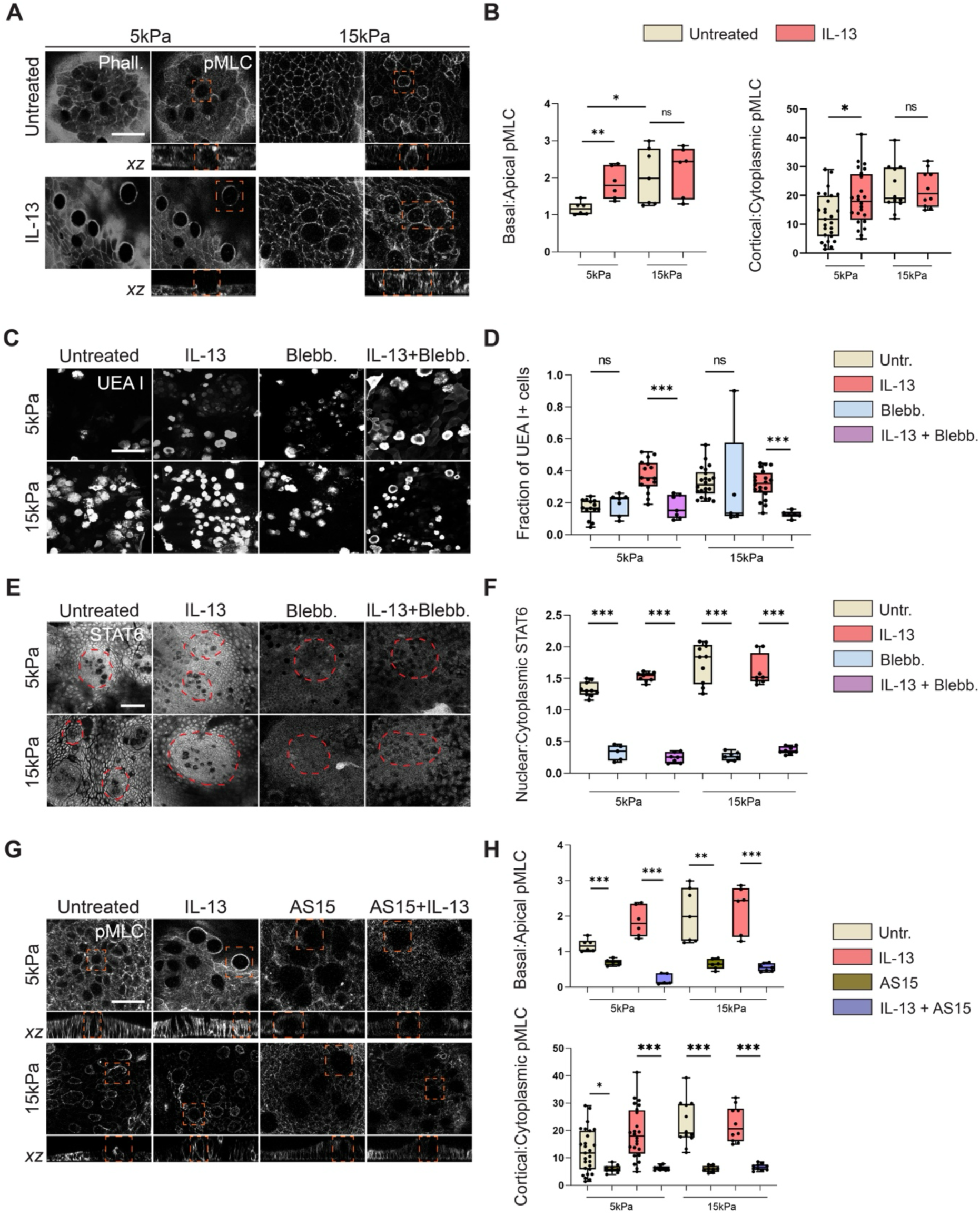
Actomyosin Contractility Links Substrate Stiffness and IL-13 to STAT6 Activation in Goblet Cells. **A.** Representative micrographs of organoid monolayers cultured on soft (0.5 kPa), intermediate (5 kPa), or stiff (15 kPa) polyacrylamide (PAA) gels, treated with or without IL-13 and immunostained for phosphorylated myosin light chain (pMLC) and stained for filamentous actin (F-actin) using fluorescently tagged phalloidin. Orange dashed boxes indicate Goblet cells in *xy* and *xz* views. SB: 20µm. **B.** Quantification of pMLC fluorescence intensity ratios of basal to apical poles and cell cortex to cytosol for conditions in *A*. For Basal:Apical quantification, each dot represents one organoid monolayer from *n* = 6, 6, 7, 6 organoid monolayers and *N* = 3, 3, 3, 3 independent experiments. For Cortical:Cytoplasm quantification, each dot represents one goblet cell from *n* = 27, 24, 12, 10 goblet cells from 6, 6, 7, 6 organoid monolayers and *N* = 3, 3, 3, 3 independent experiments. **C.** Representative micrographs of organoid monolayers cultured on intermediate or stiff PAA gels, treated with IL-13 and/or Blebbistatin (Blebb.) and stained with fluorescently tagged UEA I to identify goblet cells. Images are max projections of z-stacks. SB: 20µm. **D.** Quantification of UEA I+ secretory cell fraction for conditions in *C*. Each dot represents one organoid monolayer from *n* = 14, 7, 16, 7, 22, 5, 20, 6 organoid monolayers and *N* = 3, 3, 3, 3, 3, 3, 3, 3 independent experiments. The untreated and IL-13 data are pooled from the same data plotted in *1B*. **E.** Representative micrographs of organoid monolayers cultured on intermediate or stiff PAA gels, treated with IL-13 and/or Blebb. and immunostained for total STAT6. Red dashed lines indicate the crypt-like compartments. SB: 20µm. **F.** Quantification of nuclear to cytoplasmic STAT6 fluorescence intensity ratio for conditions in *E*. Each dot represents one organoid monolayer from *n* = 10, 5, 9, 7, 10, 7, 8, 8 organoid monolayers and *N* = 3, 3, 3, 3, 3, 3, 3, 3 independent experiments. The untreated and IL-13 data are pooled from the same data plotted in *1F*. **G.** Representative micrographs of organoid monolayers cultured on intermediate or stiff PAA gels, treated with IL-13 and/or the STAT6 inhibitor AS1517499 (AS15), immunostained for pMLC and stained for F-actin using fluorescently tagged phalloidin. Orange dashed boxes indicate corresponding Goblet cells in *xy* and *xz* views. SB: 20µm. **H.** Quantification of pMLC fluorescence intensity ratios of basal to apical poles and cell cortex to cytosol for conditions in *G*. The untreated and IL-13 data are pooled from the same data plotted in *2B.* For Basal:Apical quantification, each dot represents one organoid monolayer from *n* = 6, 6, 6, 5, 7, 6, 6, 7 organoid monolayers and *N* = 3, 3, 3, 3, 3, 3, 3, 3 independent experiments. For Cortical:Cytoplasm quantification, each dot represents one goblet cell from *n* = 27, 9, 24, 9, 12, 8, 10, 10 goblet cells from 6, 5, 6, 5, 7, 6, 6, 7 organoid monolayers and *N* = 3, 3, 3, 3, 3, 3, 3, 3 independent experiments. For panels B, D, F, H, One-way ANOVA was performed (p = 0.0248, 0.0010, 1.4e-7, 4.4e-32, 1.8e-11, 6.26e-12). Pairwise tests performed using Tukey’s HSD post-hoc text (*p<0.05, **p<0.01, ***p<0.001, ns:p>0.05).

To determine whether actomyosin contractility is required for mechanosensing- and IL-13-mediated secretory cell differentiation, we inhibited Myosin-2 activity using the specific inhibitor Blebbistatin. Inhibition of Myosin-2 rescued excess secretory cell differentiation on intermediate stiffness treated with IL-13 (Figure 2C,D), suggesting a functional interplay between cytokine signaling and mechanical force generation. Treatment with Blebbistatin on stiff substrates in the absence of IL-13 similarly restored secretory cell frequency to levels observed on soft and intermediate stiffness substrates in the absence of IL-13. These results demonstrate that myosin-mediated contractile forces are essential for transducing both inflammatory (IL-13) and biophysical cues into changes in epithelial cell differentiation.

We hypothesized that Myosin-2-mediated contractile forces could stimulate STAT6 phosphorylation and nuclear translocation, thereby linking cytoskeletal dynamics to transcriptional responses. To test this, we immunostained for STAT6 in organoid monolayers on different stiffness substrates with combined treatment of IL-13 and Blebbistatin. On intermediate stiffness, Blebbistatin treatment markedly reduced nuclear localization of total STAT6 in organoid monolayers following IL-13 treatment (Figure 2E,F). On stiff substrates, Blebbistatin treatment reduced nuclear localization of total STAT6 in organoid monolayers regardless of IL-13 treatment. These findings indicate that Myosin-driven contractility is required for the activation and nuclear accumulation of STAT6 in both cytokine-stimulated and stiffness-induced contexts. Collectively, these data support a model in which actomyosin contractility functions upstream of STAT6 activation, thereby providing a mechanistic link between microenvironment stiffness, cytoskeletal tension and cytokine-responsive transcriptional regulation in the intestinal epithelium.

To investigate whether STAT6 signaling in turn regulates actomyosin contractility, we treated organoid monolayers on intermediate stiffness with IL-13 in the presence or absence of AS15 and investigated pMLC localization. Inhibition of STAT6 significantly reduced cortical pMLC localization, suggesting that STAT6 activity may facilitate assembly or stabilization of actomyosin structures in response to IL-13 signaling (Figure 2G). Intriguingly, AS15 treatment also reduced cortical pMLC accumulation in organoids cultured on stiff substrates without IL-13, indicating that STAT6 may contribute to stiffness-induced cytoskeletal remodeling (Figure 2G,H). These findings point to a potential feedback loop in which STAT6 not only responds to mechanical and biochemical signals but may also modulate cytoskeletal architecture under inflammatory or fibrotic conditions.

### Substrate mechanosensing and IL-13 signaling modulate cell-cell and cell-substrate adhesion integrity

Previous studies have demonstrated that focal adhesion formation and cellular mechanical force generation are significantly influenced by substrate stiffness.

However, the role of IL-13 signaling in modulating cellular force generation and FA formation and maturation remains poorly understood. We therefore investigated the effects of substrate stiffness and IL-13 signaling using traction force microscopy (TFM). Organoid monolayers cultured on intermediate (5 kPa) substrates exhibited distinct spatial patterns of stress magnitude (Figure 3A). In untreated conditions, traction stress magnitude was approximately 20 Pa at the crypt center and gradually increased toward the villus region, reaching peak values of approximately 60 Pa (Figure 3B). This spatial gradient reflects the mechanical compartmentalization characteristic of intestinal organoid architecture^31^. Following IL-13 treatment of organoid monolayers on intermediate stiffness substrates, stress magnitude at the crypt center increased to approximately 30 Pa, significantly higher compared to untreated conditions. This increase in mechanical stress generation specifically within the stem cell compartment suggests that IL-13 enhances the contractile activity of intestinal stem cells, potentially through modulation of actomyosin contractility pathways. The elevated stress at the crypt center may contribute to enhanced mechanotransduction signaling critical for stem cell fate decisions. Conversely, in the villus region, IL-13 treatment resulted in approximately 30% lower stress magnitude compared to untreated monolayers, indicating that IL-13 exerts opposing mechanical effects in the crypt-like and villus-like compartments. Splitting the traction stresses into radial and tangential components with respect to the contour of the crypt-like region, we found that in organoid monolayers treated with IL-13, traction stresses in the crypt center were outward facing as opposed to inward facing tractions in untreated monolayers (Figure 3B). Organoid monolayers on stiff substrates demonstrated markedly higher stress magnitudes at the crypt center compared to intermediate stiffness substrates, confirming that increased substrate stiffness promotes enhanced traction force generation as shown previously in other cellular models^48–50^ (Figure 3C). Notably, IL-13 treatment on stiff substrates did not further alter stress magnitude in the crypt, suggesting that the effects of IL-13 on traction stress production may be stiffness-dependent or that contractile machinery becomes saturated at higher substrate rigidities.

**Figure 3.**
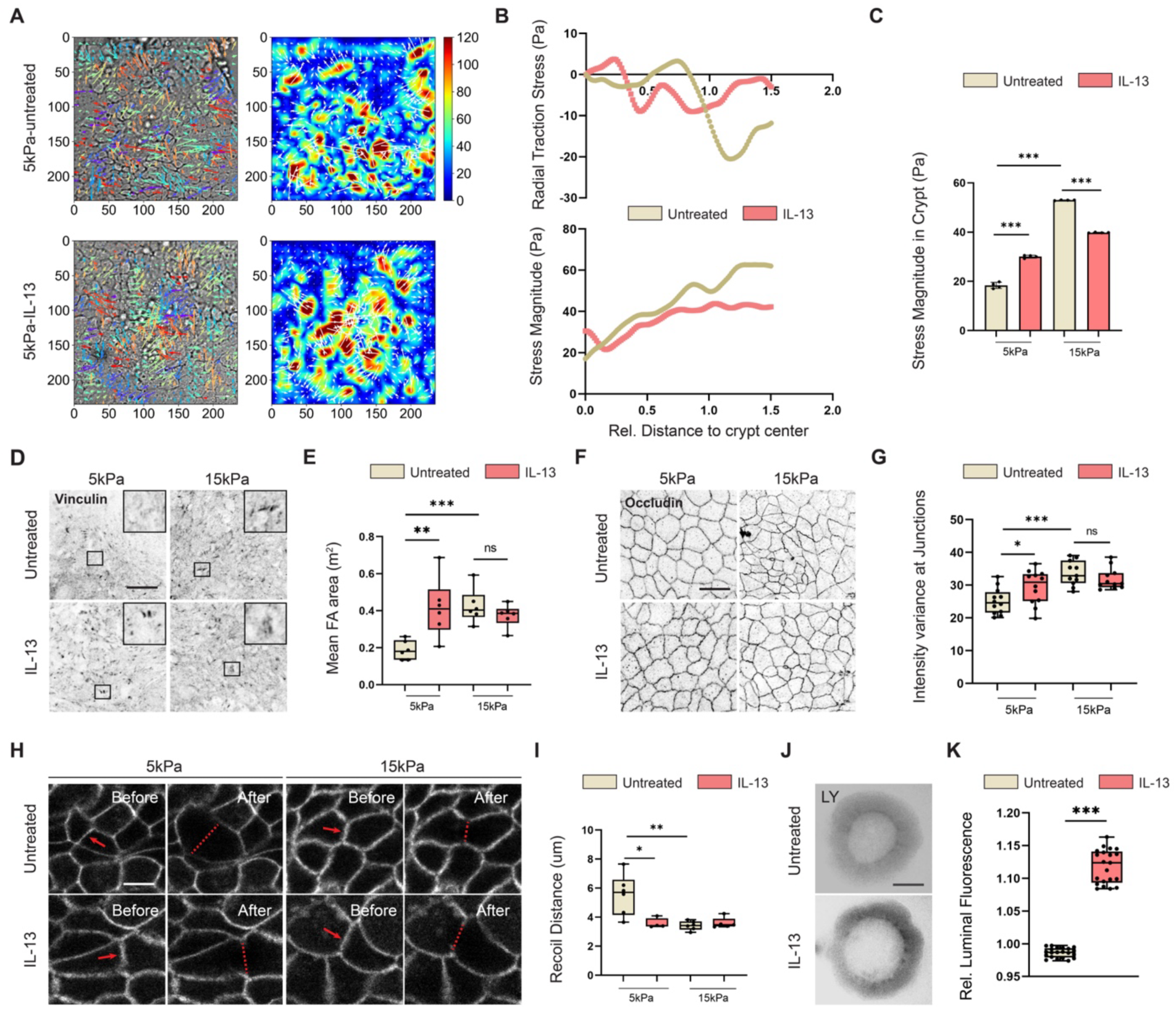
Substrate mechanosensing and IL-13 signaling modulate cell-cell and cell-substrate adhesion integrity. **A.** Representative traction force microscopy (TFM) maps of organoid monolayers cultured on intermediate (5 kPa) polyacrylamide (PAA) gels treated with or without IL-13. **B.** Quantification of mean traction force magnitude and radial stress magnitude exerted by organoid monolayers cultured on intermediate (5kPa) or stiff (15kPa) PAA gels. Relative distances to the crypt are rescaled such that the crypt center is at position 0, the crypt edge at position 1, and the villus-like compartment at values >1. **C.** Bar plot representing mean ± SEM of stress magnitude within the crypt region (averaged from rescaled position 0-1. Each dot represents one organoid monolayer from *n* = 4, 4, 4, 4 and *N* = 2, 2, 2, 2 independent experiments. **D.** Representative micrographs of organoid monolayers cultured on intermediate or stiff PAA gels, treated with or without IL-13 and immunostained for the focal adhesion protein vinculin. Black squares indicate zoomed inset region. SB: 20µm. **E.** Quantification of total focal adhesion (FA) area per field of view for conditions in *C*. Each dot in the box plot represents one organoid monolayer from *n* = 6, 6, 6, 6 and *N* = 3, 3, 3, 3 independent experiments. **F.** Representative micrographs of organoid monolayers cultured on intermediate/stiff PAA gels, treated with or without IL-13 and immunostained for the tight junction protein occludin. SB: 20µm. **G.** Quantification of variance in occludin signal density along individual junctions from individual cell segmentation for conditions in *E*. Variance was normalized to the mean fluorescence intensity of each respective junction and averaged across all junctions per image. Each dot in the box plot represents one organoid monolayer from *n* = 12, 12, 11, 11 and *N* = 3, 3, 3, 3 independent experiments. **H.** Representative micrographs from two-photon laser ablation of mT/mG organoid monolayers on intermediate/stiff PAA gels, with or without IL-13, before and after ablation of individual cell-cell junctions. Red arrow indicates the cell junction that was ablated. Red dashed line indicates distance between two vertices of the junctions after the laser ablation. SB: 10µm. See also Supplementary Video 2. **I.** Quantification of junctional recoil distance (μm) immediately following two-photon laser ablation of cell-cell junctions for conditions in *G*. Each dot in the box plot represents one organoid monolayer from *n* = 6, 4, 6, 5 and *N* = 3, 3, 3, 3 independent experiments. **J.** Representative micrographs of 3D intestinal organoid with or without IL-13 pre-treatment, 5 h after adding Lucifer Yellow (LY) to the organoid medium. SB: 100µm. **K.** Quantification of relative luminal fluorescence (organoid lumen to medium fluorescence intensity) over time after adding LY. Each dot in the box plot represents one organoid from *n* = 21, 21 and *N* = 3, 3 independent experiments. For panels *C, E, G, I, K*, One-way ANOVA was performed (p = 3.0e-16, 0.0017, 6.6e-5, 0.0010, 2.3e-24). Pairwise tests performed using Tukey’s HSD post-hoc text (*p<0.05, **p<0.01, ***p<0.001, ns:p>0.05).

Increased traction forces are associated with increased formation and maturation of cell-substrate adhesions^51^. Thus, we next investigated the distribution of vinculin, a key mechanosensitive component of FAs. To this end, we cultured organoid monolayers on different stiffness substrates, treated with IL-13 and immunostained for vinculin. Treatment with IL-13 on intermediate stiffness substrates led to increased formation of vinculin-positive FAs, as indicated by increased FA area (Figure 3D, E). FA area was also higher on stiff substrates compared to intermediate substrates, and IL-13 treatment on stiff substrates had no further effect on FA area. In addition to FA area, the relative intensity of vinculin in FAs was also increased for IL-13 treatment on intermediate stiffness substrates or on stiff substrates regardless of IL-13 treatment (Supplementary Figure 4A). In contrast, the relative intensity of paxillin and phosphorylated focal adhesion kinase (pFAK) in FAs did not change for different substrate stiffnesses with or without IL-13 treatment (Supplementary Figure 4B,C). The difference in staining between vinculin and paxillin/pFAK could be explained by the fact that vinculin recruitment is force-dependent^52^. Paxillin and pFAK, on the other hand, mark early assembly and kinase activity at FAs and could already be near-saturated in polarized epithelia on ECM, yielding limited dynamic range to measure changes in paxillin or pFAK accumulation under higher mechanical stresses at FAs. Together, these data suggests that substrate stiffness and cytokine treatment not only modulate traction stress generation, but also influence FA formation and maturation.

As FA area increases, cells become more strongly anchored to the substrate, which physically restricts their ability to move relative to one another. This results in reduced collective cell movement and can influence jamming/unjamming transitions in epithelial monolayers^53–55^. To test this in the context of mechanosensing and IL-13 signaling, we measured epithelial monolayer fluidity using particle imaging velocimetry (PIV) on monolayers cultured on intermediate and stiff substrates. On intermediate stiffness, cells exhibited strong spatial differences in mean instantaneous speed, with fast-moving cells typically localized to crypt regions (Supplementary Figure 4D, Supplementary Video 1). In contrast, monolayers on stiff substrates displayed a significant decrease in migration speed, suggesting that increased focal adhesion strength on stiffer substrates indeed restricts collective cell movement within the monolayer (Supplementary Figure 4E). Inhibition of STAT6 using AS15 led to a significant increase in migration speed compared to untreated controls on stiff substrates, suggesting that STAT6 signaling is involved in the reduction of mechanosensing-mediated migration.

Epithelial integrity depends on a balance between cell-matrix adhesions and cell-cell adhesions^56^. This mechanical interplay between adhesion types suggests that enhanced substrate engagement through integrin-based adhesions could come at the expense of AJ- and TJ-mediated cohesion^57,58^. Increased FA formation and traction force generation on stiffer substrates or in the presence of IL-13 could therefore mechanically compete with or destabilize cell-cell junctions, leading to compromised barrier function^59,60^. To address this, we cultured organoid monolayers on different stiffness substrates with or without IL-13 treatment and immunostained for the TJ protein occludin. Organoid monolayers cultured on intermediate stiffness substrates displayed well-defined TJs localized at apical cell interfaces (Figure 3F). Monolayers cultured on intermediate stiffness substrates with IL-13 and monolayers cultured on stiff substrates displayed higher junctional intensity variance, indicating more discontinuous and irregular junctions (Figure 3G). In parallel, we investigated the TJ scaffolding protein Zonula Occludins-1 (ZO-1). On untreated monolayers on intermediate stiffness, ZO-1 was localized in an apically polarized manner (Supplementary Figure 4F). Upon treatment with IL-13, relative ZO-1 intensity at junctions was significantly reduced (Supplementary Figure 4G). On stiff substrates, relative junctional ZO-1 intensity was also reduced as compared to intermediate stiffness, and further addition of IL-13 on stiff substrate had no additional effect. These data suggest that both IL-13 and high substrate stiffness disrupt tight junction morphology, likely reflecting reduced junctional integrity.

As cell-cell junction complex assembly is a mechanosensitive process, the observed disruption of TJs associated with increased substrate stiffness and IL-13 signaling could reflect a difference in mechanical tension at cell-cell junctions. We therefore measured relative junctional tension using two-photon laser ablation of individual cell-cell junctions in organoid monolayers on different stiffness substrates in the absence or presence of IL-13 (Figure 3H, Supplementary Video 2). As a proxy for junctional tension, we quantified junctional recoil distance immediately following ablation. Junctional recoil was reduced in IL-13-treated monolayers on intermediate stiffness substrates, indicating lower junctional tension (Figure 3I). Similarly, on stiff substrates, junctional recoil was significantly lower compared to monolayers cultured on intermediate stiffness substrates. IL-13 addition on stiff substrates had no additional effect on junctional tension. Taken together, these experiments indicate that high substrate stiffness and IL-13 enhance cell-substrate adhesions by increasing traction forces and FA formation. However, this comes at the cost of disruptions in cell-cell adhesions, as suggested by aberrant tight junction structure and reduced junctional tension.

Weakened cell-cell adhesion could give rise to disruptions in barrier function, characterized by increased epithelial permeability. To directly assess how these changes impact barrier function, we experimentally measured organoid permeability using a previously described assay^61^. To this end, we pre-treated freshly passaged 3D organoids with IL-13 and subsequently supplemented the medium with Lucifer Yellow (LY), a small fluorophore that can diffuse through the pericellular space when epithelial barrier is disrupted. We then performed live fluorescence microscopy and quantified the relative LY fluorescence intensity in the organoid lumen over time (Fig. 3J,K). While LY intensity in control organoids remained stable over time, the relative luminal intensity in IL-13-treated organoids gradually increased. This resulted in significantly higher relative luminal intensity after 5 hours for IL-13-treated organoids, suggesting that IL-13 treatment disrupts barrier integrity. These results support the conclusion that alterations in cell-cell junctions and adhesions induced by IL-13 has functional consequences for epithelial barrier function.

### Substrate Stiffness and IL-13 Synergistically Regulate YAP Nuclear Translocation via Actomyosin Contractility

We next sought to elucidate the downstream signaling mechanisms mediating mechanosensing and IL-13 signaling. To this end, we first investigated the localization and activation of YAP, a key mechanosensory protein previously implicated in intestinal stem cell maintenance and differentiation^37^. We cultured 2D organoid monolayers on different stiffness substrates, treated with IL-13 and immunostained for YAP. For organoid monolayers on soft substrates (0.5 kPa), YAP levels were generally low, with diffuse localization patterns and low relative nuclear YAP (Figure 4A,B). Upon IL-13 treatment, we observed no significant change in relative nuclear YAP localization. On intermediate stiffness (5 kPa), YAP expression was restricted to differentiated cells in villus-like regions, where cells displayed enriched nuclear YAP. With IL-13 treatment on intermediate substrates, nuclear YAP increased in the crypt-like compartment. Similarly, organoid monolayers on stiff substrates (15 kPa) with or without IL-13 treatment displayed increased total and relative nuclear YAP in the crypt-like region. To determine whether changes in nuclear YAP localization are also associated with changes in overall YAP expression, we harvested cell lysates from the organoid monolayers and performed Western blot analysis. YAP expression increased slightly following IL-13 treatment for monolayers on intermediate and stiff substrates, though there was no difference in baseline expression for untreated monolayers on intermediate or stiff substrates (Fig. 4C,D). Together, these data suggest that both increased substrate stiffness and IL-13 can drive YAP nuclear localization, while only IL-13 influences YAP expression.

**Figure 4.**
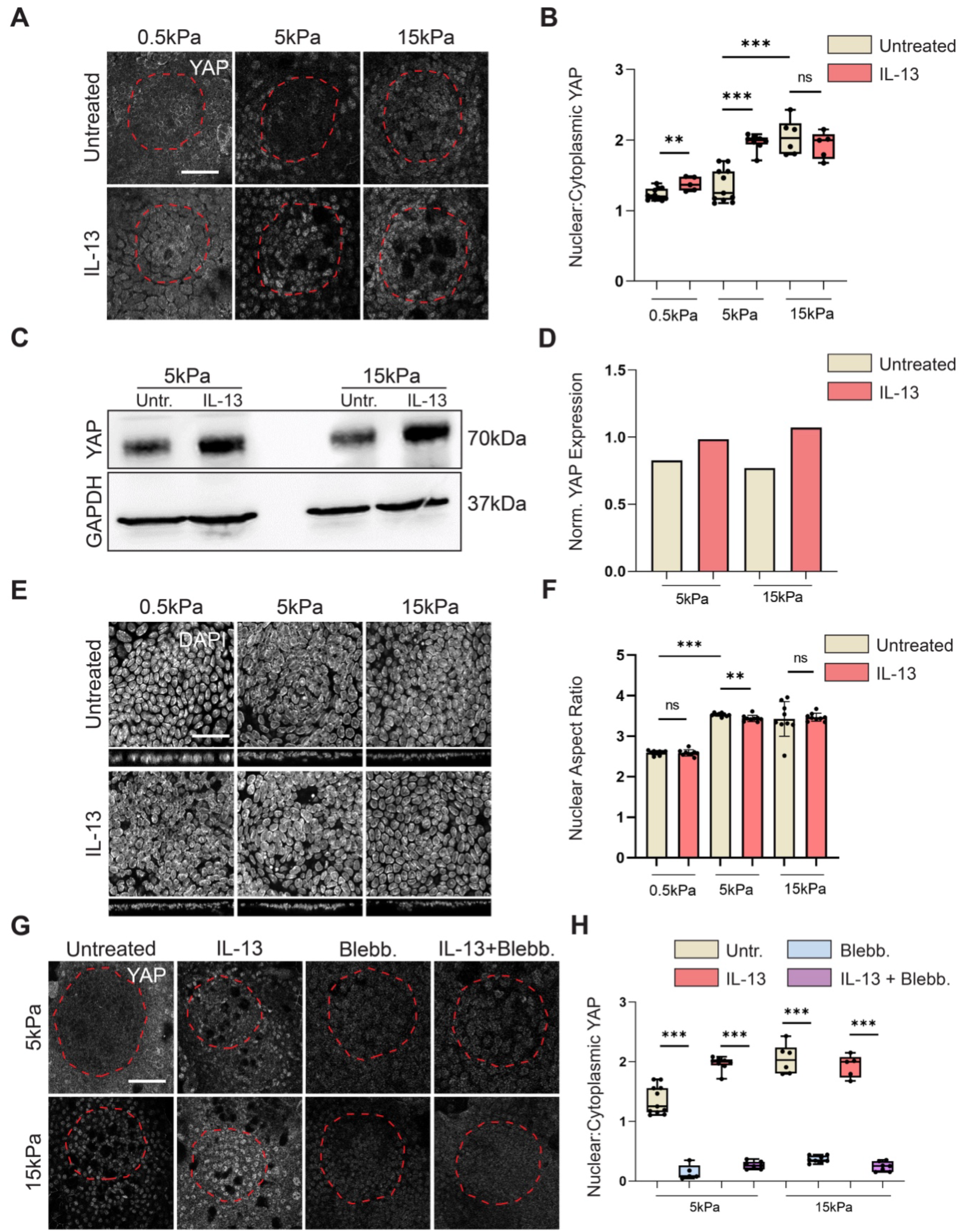
Substrate Stiffness and IL-13 Synergistically Regulate YAP Nuclear Translocation via Actomyosin Contractility. **A.** Representative micrographs of organoid monolayers cultured on soft (0.5 kPa), intermediate (5 kPa), or stiff (15 kPa) polyacrylamide (PAA) gels, treated with or without IL-13 and immunostained for YAP. Dashed red lines indicate the border of the crypt-like regions. Red dashed lines indicate the crypt-like compartments. SB: 20µm. **B.** Quantification of nuclear-to-cytoplasmic YAP fluorescence intensity ratio for conditions in *A*. Each dot represents one organoid monolayer from *n* = 12, 5, 11, 8, 6, 5 organoid monolayers and *N* = 3, 3, 3, 3, 3, 3 independent experiments. **C.** Western blot of YAP expression in organoid monolayers cultured on intermediate/stiff substrates with or without IL-13 treatment. **D.** Densitometric quantification of Western blots as in *C*. Bar plot represents mean±SEM. **E.** Representative micrographs of organoid monolayers cultured on soft/intermediate/stiff PAA gels, treated with or without IL-13 and nuclear staining using DAPI for 3D segmentation and morphometric analysis. SB: 20µm. **F.** Quantification of nuclear aspect ratio (ratio of longest to shortest axis from ellipsoid fit) for cells in crypt-like regions for conditions in *E*. Each dot represents one organoid monolayer from *n* = 9, 9, 9, 9 9, 9 organoid monolayers and *N* = 3, 3, 3, 3, 3, 3 independent experiments. **G.** Representative micrographs of organoid monolayers cultured on soft/intermediate/stiff PAA gels, treated with IL-13 or the Myosin-2 inhibitor Blebbistatin (Blebb.) and immunostained for YAP. Red dashed lines indicate the crypt-like compartments. SB: 20µm. **H.** Quantification of nuclear-to-cytoplasmic YAP ratio post-Blebbistatin treatment. Each dot represents one organoid monolayer from *n* = 11, 5, 8, 7, 6, 8, 5, 7 organoid monolayers and *N* = 3, 3, 3, 3, 3, 3, 3, 3 independent experiments. The untreated and IL-13 data are pooled from the same data plotted in *4B*. For panels *B, F, H*, One-way ANOVA was performed (p = 7.2e-14, 1.8e-18, 3.1e-33). Pairwise tests performed using Tukey’s HSD post-hoc text (**p<0.01, ***p<0.001, ns:p>0.05).

Previous studies have demonstrated that nuclear YAP accumulation can be mediated by nuclear deformation, which occurs through mechanical coupling of the cytoskeleton to the nucleus via focal adhesions, the actin cytoskeleton, and the LINC complex^62,63^. We therefore hypothesized that IL-13 signaling or mechanosensing could regulate nuclear YAP accumulation by inducing nuclear deformations. To test this, we cultured organoid monolayers on intermediate or stiff substrates with or without IL-13 treatment, fixed and stained for cell nuclei and visualized by confocal microscopy (Figure 4E). Using the resulting images, we performed 3D segmentation of nuclei and quantified nuclear morphometrics of crypt cells. Specifically, we investigated changes in nuclear shape by measuring the aspect ratio (AR) of the longest to shortest axis from a 3D ellipsoid fit (Supplementary Figure 5A). For organoid monolayers cultured on intermediate or stiff substrates, the nuclear AR was significantly higher compared to soft stiffness, consistent with previous studies^64^ (Fig. 4E,F). On soft or stiff PAA gels, IL-13 treatment had no effect on nuclear AR, while on intermediate stiffness, nuclear AR decreased slightly, but significantly, upon addition of IL-13. Neither nuclear orientation nor the ratio of the smallest two principal axes (R2/R3) was influenced by substrate stiffnesses or IL-13 treatment (Supplementary Figure 5B). Together, these findings suggest that nuclear deformation could act as a mechanical sensor to regulates YAP-mediated mechanotransduction in response to substrate stiffness. However, the overall changes in nuclear morphology in intestinal epithelial cells are subtle compared with single mesenchymal cells that were analyzed in previous studies, possibly due to the overall lower traction stresses produced by polarized epithelial cells^65^.

Our results suggest that IL-13 treatment promotes increased cortical pMLC localization, and inhibition of Myosin-2 activity rescues goblet cell hyperplasia following IL-13 treatment (Fig. 2A-D). As these results suggest that actomyosin contractility is crucial for secretory cell differentiation, and given that YAP nuclear localization is known to be regulated by cytoskeletal tension, we next tested whether actomyosin contractility also controls YAP nuclear translocation in this context. To this end, we cultured organoid monolayers on substrates of varying stiffness PAA gels and treated with IL-13 or Blebbistatin, followed by immunostaining for YAP. On intermediate stiffness substrates, Blebbistatin treatment markedly reduced the relative nuclear localization of YAP after IL-13 treatment (Fig. 4G,H). Notably, Blebbistatin also reduced nuclear YAP localization on stiff substrates even without IL-13 treatment. These findings indicate that myosin-driven contractility is required for the activation and nuclear accumulation of YAP driven by both cytokine stimulation and increased substrate stiffness. Collectively, these data support a model in which increased actomyosin contractility in response to IL-13 or high substrate stiffness promotes nuclear translocation of YAP.

### Substrate Stiffness and IL-13 Promote Secretory Cell Differentiation via STAT6- Dependent YAP Activation

To directly assess the functional role of YAP in secretory cell differentiation driven by IL-13 and mechanosensing, we cultured 2D organoid monolayers on intermediate and stiff substrates and treated with IL-13 in the presence or absence of the YAP inhibitor Verteporfin. Labeling of goblet cells using fluorescently tagged UEA I revealed that for organoid monolayers on intermediate or stiff substrates treated with IL-13, Verteporfin restored secretory cell frequency to levels similar to untreated monolayers on intermediate substrates (Figure 5A,B). Interestingly, Verteporfin also rescued goblet cell hyperplasia for monolayers on stiff substrates in the absence of IL-13. These results indicate that YAP activation is required for increased secretory cell differentiation stimulated by mechanosensing or IL-13 signaling, and that inhibition of YAP can reverse this effect in both contexts.

**Figure 5.**
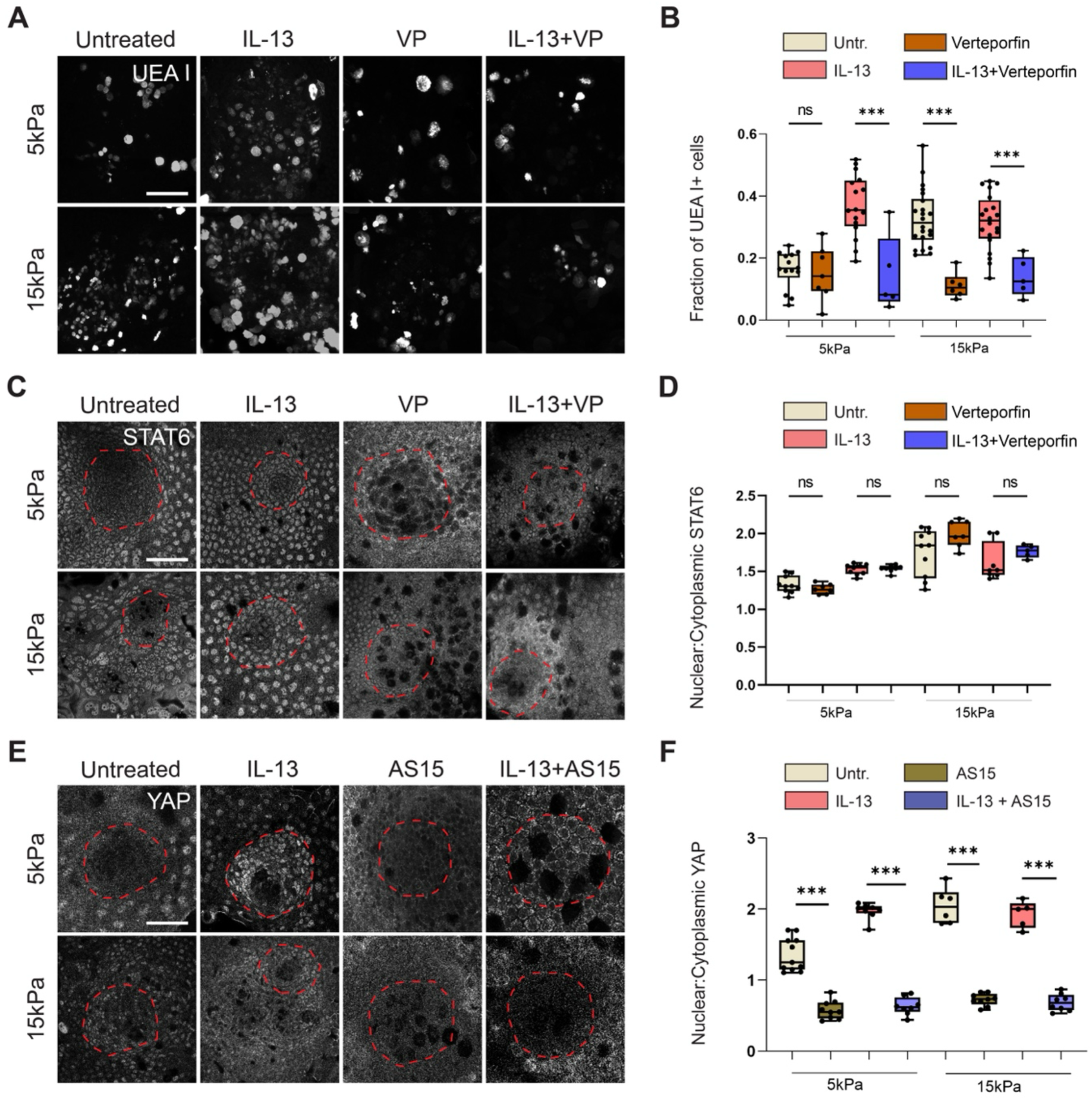
Substrate Stiffness and IL-13 Promote Secretory Cell Differentiation via STAT6-Dependent YAP Activation. **A.** Representative micrographs of organoid monolayers cultured on intermediate (5 kPa) or stiff (15 kPa) polyacrylamide (PAA) gels, treated with IL-13 or the YAP inhibitor Verteporfin (VP) and stained with fluorescently labeled UEA I to label secretory cells. Images are max projections of z-stacks. SB: 20µm. **B.** Quantification of the fraction of UEA I+ secretory cells for conditions as in *A*. Each dot represents one organoid monolayer from *n* = 14, 7, 16, 5, 22, 6, 20, 5 organoid monolayers and *N* = 3, 3, 3, 3, 3, 3, 3, 3 independent experiments. The untreated and IL-13 data are pooled from the same data plotted in *1B*. **C.** Representative micrographs of organoid monolayers cultured on intermediate/stiff PAA gels, treated with IL-13 or the YAP inhibitor Verteporfin (VP) and immunostained for STAT6. Red dashed lines indicate the crypt-like compartments. SB: 20µm. **D.** Quantification of nuclear-to-cytoplasmic STAT6 fluorescence intensity ratios for conditions in *C*. Each dot represents one organoid monolayer from *n* = 10, 7, 9, 8, 10, 7, 8, 5 organoid monolayers and *N* = 3, 3, 3, 2, 3, 3, 3, 2 independent experiments. The untreated and IL-13 data are pooled from the same data plotted in *1F*. **E.** Representative micrographs of organoid monolayers cultured on intermediate/stiff PAA gels, treated with IL-13 or the STAT6 inhibitor AS1517499 (AS15) and immunostained for YAP. Red dashed lines indicate the crypt-like compartments. SB: 20µm. **F.** Quantification of nuclear-to-cytoplasmic YAP fluorescence intensity ratios for conditions in *E*. Each dot represents one organoid monolayer from *n* = 11, 9, 8, 8, 6, 9, 5, 8 organoid monolayers and *N* = 3, 2, 3, 3, 3, 3, 3, 2 independent experiments. The untreated and IL-13 data are pooled from the same data plotted in *4B*. For panels *B, D*, *F*, One-way ANOVA was performed (p = 3.7e-13, 1.5e-10, 6.3e-31). Pairwise tests performed using Tukey’s HSD post-hoc text (***p<0.001, ns:p>0.05).

Our results suggest that both STAT6 and YAP are essential mediators of IL-13 signaling and mechanosensing. However, it is not clear whether STAT6 and YAP are activated in parallel, or whether their activation is co-dependent. To address this, we treated organoid monolayers cultured on intermediate and stiff substrates and treated with IL-13 and Verteporfin. Verteporfin did not significantly alter nuclear localization of STAT6 on any of the tested substrate stiffnesses, even in the presence of IL-13 (Figure 5C,D). This indicates that STAT6 activation is independent and potentially upstream of YAP in driving increased secretory cell differentiation. To investigate whether STAT6 acts upstream of YAP, we inhibited STAT6 using AS15 and assessed relative YAP nuclear localization. On intermediate substrates, AS15 treatment reduced YAP nuclear localization in response to IL-13 (Figure 5E,F). AS15 treatment similarly reduced relative nuclear YAP localization in organoid monolayers on stiff substrates, either with or without IL-13 treatment. These results suggest that STAT6 functions upstream to promote nuclear accumulation of YAP.

Together, these data support a model in which IL-13 and high substrate stiffness promote secretory cell differentiation through STAT6-dependent YAP nuclear translocation and activation, which in turn drives secretory lineage specification (Figure 6). Notably, our findings suggest that, in contrast to previous studies using single mesenchymal cells, YAP nuclear localization during mechanosensing in polarized epithelial cells is not solely a direct consequence of mechanical cues, but also requires upstream activation by STAT6.

**Figure 6.**
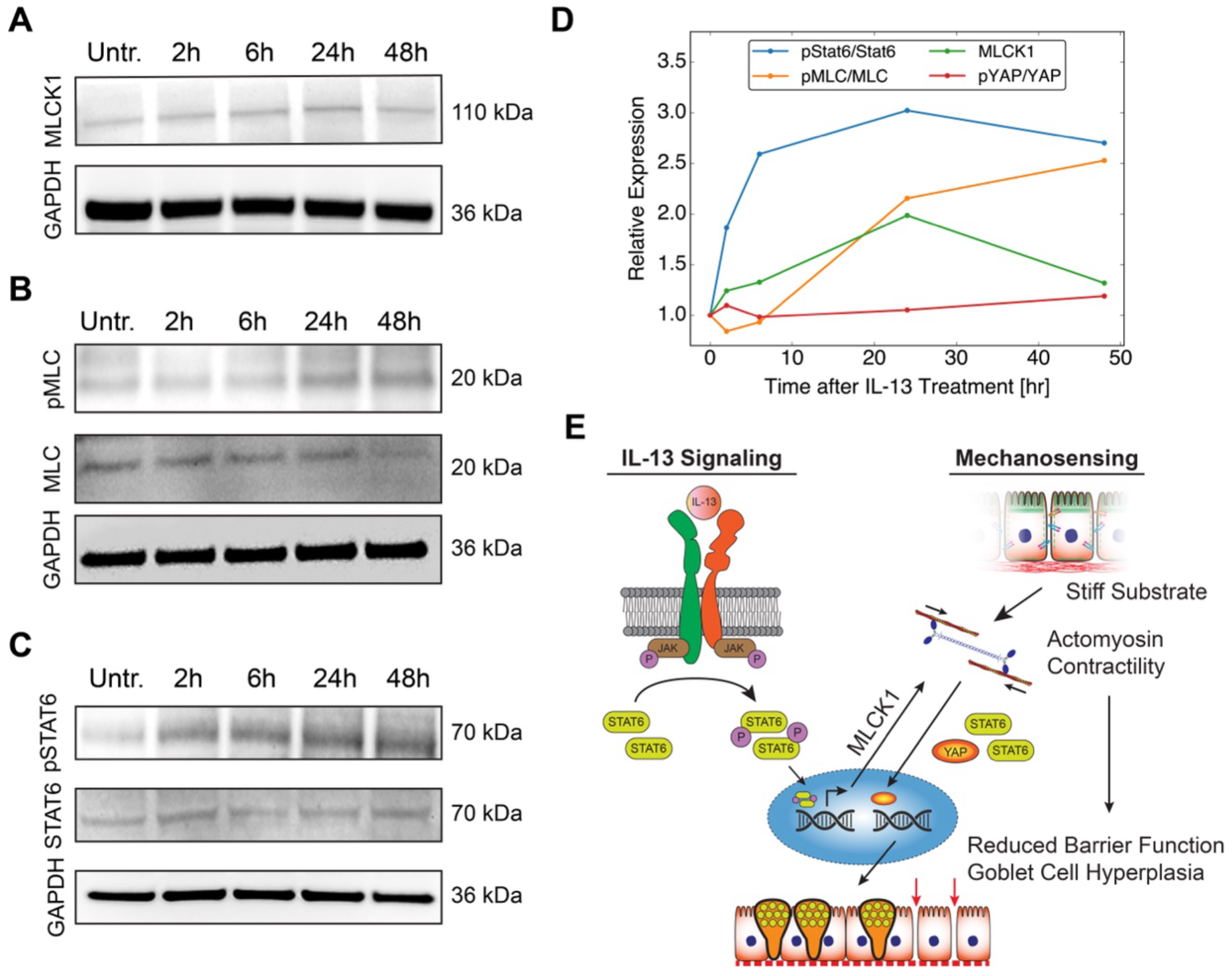
Mechanosensing and IL-13 Signaling Synergistically Modulate Intestinal Stem Cell Differentiation via STAT6, YAP and Actomyosin Contractility. A-C. Western blots of 3D organoid lysates after different timepoints following IL-13 treatment or without IL-13 treatment (Untr.). Lysates were blotted for total MLCK1 expression (*A*) and total or phosphorylated YAP (*B*) and STAT6 (*C*). **D.** Quantification of relative expression and phosphorylation of STAT6, MLCK1, MLC and YAP at different timepoints following IL-13 treatment or without IL-13 treatment (t=0hr). **E.** Schematic representation of mechanism underlying synergies and feedback between IL-13 signaling and mechanosensing in regulating intestinal epithelial differentiation and barrier function.

### STAT6 Promotes Actomyosin Contractility and YAP Activation via MLCK1 Expression

The observation that YAP nuclear localization requires upstream STAT6 activation prompts the question of how STAT6 and YAP are mechanistically linked. Our data indicated that IL-13 treatment increased traction force generation and focal adhesion formation and that IL-13-mediated YAP nuclear localization requires actomyosin contractility (Fig. 3A-C, Figure 4G,H). We therefore hypothesized that STAT6 could increase contractility, thereby driving YAP mechanotransduction. Previous work has identified MLCK1 as a downstream target of STAT6 in intestinal epithelial cells *in vivo*^66^. To determine how STAT6 activation modulates MLCK1 expression and activity, we treated 3D intestinal organoids with IL-13 and harvested lysates at 2h, 6h, 24h and 48h after treatment. We then performed Western blot analysis to determine STAT6 phosphorylation, MLCK1 expression and MLC phosphorylation as an indicator of active contractility (Figure 6A-C). IL-13 treatment induced a rapid increase in STAT6 phosphorylation that reached a plateau after 6h (Figure 6D). MLCK expression gradually increased after IL-13 treatment, reaching a maximum at 24h, while MLC phosphorylation was slightly delayed, beginning to increase 6h after IL-13 treatment and continuing to increase 48h after treatment. This corresponds with the increase in nuclear YAP accumulation in IL-13-treated organoid monolayers, which depends on actomyosin contractility, suggesting that contractility mechanistically links STAT6 and YAP. To test whether STAT6 and YAP could directly interact, we performed a co-immunoprecipitation using STAT6 as bait. Western blot analysis indeed indicated a direct interaction between STAT6 and YAP, which was slightly enhanced following IL-13 treatment (Supplementary Figure 6A). Together, these data suggest that in addition to increasing contractility and nuclear deformation via STAT6-dependent expression of MLCK1, STAT6 and YAP could be cooperatively imported into the nucleus via direct biochemical interaction.

A possible alternative mechanism by which STAT6 could modulate YAP activity is through Hippo signaling, which regulates YAP nuclear localization via phosphorylation. Indeed, prior work has identified links between cytokine signaling and Hippo pathway^67^. To first establish whether the Hippo pathway is active in intestinal organoids, we treated 3D organoids with NIBR-LTSi, an inhibitor of the upstream Hippo kinase large tumor suppressor (LATS)^68^. Western blot analysis of organoid lysates suggested that NIBR-LTSi dramatically reduced levels of pYAP both with or without IL-13 treatment (Supplementary Figure 6B). However, we did not observe any significant change in YAP phosphorylation in response to IL-13 at any of the tested timepoints (Figure 6D, Supplementary Figure 6C). These data suggest that while Hippo signaling is active during normal intestinal epithelial homeostasis, the Hippo pathway is not involved in the IL-13 response. We therefore conclude that YAP activation in response to IL-13 is mediated by STAT6-dependent MLCK1 expression, with a potential additional contribution by direct STAT6-YAP binding (Figure 6E).

## Discussion

Goblet cell hyperplasia is a hallmark of chronic intestinal inflammation, often linked to activation of type 2 cytokines such as IL-13^18^. Chronic inflammation in the intestine and other tissues can also lead to long-term complications including increased tissue stiffness and fibrosis due to excess ECM deposition and remodeling. However, the influence of biomechanical cues, and particularly substrate stiffness, on epithelial cell fate and their cross-talk with inflammatory cytokine signaling pathways remains poorly understood. In this study, we uncover a synergistic interplay between mechanosensing and IL-13 signaling in promoting goblet cell differentiation in the intestinal epithelium via a STAT6- and YAP-dependent mechanism. This convergence of mechanical and cytokine signaling provides novel insight into how epithelial homeostasis is perturbed in pathological settings and suggests potential therapeutic strategies for targeting mechano-inflammatory crosstalk in diseases such as IBDs.

We found that intermediate stiffness substrates (5 kPa) mechanically primed intestinal epithelial cells to respond to IL-13 treatment. The optimization of signaling at 5 kPa also coincides with 5 kPa being a critical threshold stiffness in other cellular systems for regulating cell morphology and organization. The force magnitudes required to produce deformations of substrate stiffnesses of 5 kPa corresponds with the force production of molecular motors, and this value has been proposed to serve as an internal stiffness reference for cellular mechanosensing^43^. Our data align with prior studies in lung epithelium and fibroblasts, where substrate stiffness enhances responsiveness to TGF-β and other cytokines, suggesting a conserved paradigm in tissue remodeling and inflammation^48^. Strikingly, stiff substrates (15 kPa) promoted secretory cell hyperplasia even in the absence of IL-13. These findings suggest that epithelial cells interpret stiff environments as inflammatory-like cues, independent of canonical immune stimuli.

Mechanistically, increasing substrate stiffness augmented cortical pMLC localization and traction force generation, and IL-13 amplified these responses exclusively at intermediate stiffness, reinforcing the idea of stiffness-mediated priming (Figure 2A,B; Figure 3A-C). Subcellular vinculin localization confirmed that focal adhesion area increased at intermediate stiffness, with IL-13 exerting no further effect on already stiff substrates (Figure 3D,E). However, this mechanical reinforcement came at the cost of epithelial cohesion: IL-13 and high substrate stiffness disrupted tight junction architecture and reduced junctional tension, highlighting a trade-off between enhanced cell-matrix adhesion and loss of cell-cell contacts (Figure 3F-I). This mechanical imbalance also compromised barrier function in organoids (Figure 3J,K), suggesting that fibrosis in advanced IBD could also increase barrier permeability leading to “leaky gut,” a condition often associated with intestinal inflammation^69^. Our results suggest that leaky gut can be caused not only via transcriptional regulation of junctional proteins but also via mechanically mediated junction destabilization^70^. Furthermore, this finding opens new questions and avenues of research focused on mechanisms regulating the balance between adhesion types in epithelial tissues.

Central to our findings is the identification of STAT6 as a key integrator of mechanical and cytokine signaling. Our data on nuclear morphometrics indicate that STAT6 nuclear localization on intermediate or stiff substrates could be induced by nuclear deformation (Figure 4E,F). Based on previous studies, we predict that this change in nuclear shape is a consequence of changes in mechanical tension at cell-substrate adhesions and/or cell-cell junctions that are transmitted to the nucleus via the actomyosin cytoskeleton and the LINC complex^71–73^. IL-13 induced STAT6 phosphorylation and nuclear entry on intermediate stiffness, whereas stiff substrates appeared to trigger phosphorylation-independent STAT6 nuclear accumulation (Figure 1C-F), suggesting an alternative, mechanosensitive activation mechanism. This is also supported by previous findings in macrophages linking cell and nuclear elongation to STAT6 nuclear accumulation^23^. However, this has not previously been shown in polarized epithelial cells. In the case of YAP, nuclear elongation is proposed to increase the mechanical tension of the nuclear membrane, inducing dilation of nuclear pore complexes. This modulates the relative import and export rates of mechanosensitive proteins like YAP, leading to nuclear accumulation^71,74,75^. It is thus possible that STAT6 nuclear accumulation is regulated in a similar manner. However, our TFM results highlight important differences in force generation in single cells vs. epithelial cell sheets (Figure 3A-C). In organoid monolayers, the magnitudes of in-plane traction stresses (up to ∼100 Pa), are substantially lower than traction stresses produced by single cultured cells like fibroblasts (∼800-1200 Pa). The traction forces required to produce nuclear deformation and nuclear YAP accumulation in single cells are over ∼1000 Pa, an entire order of magnitude larger than the maximal stresses produced by polarized epithelial monolayers^31,76^. Furthermore, in single adherent cells, nuclei are typically oriented parallel with the substrate, while our finding suggest that nuclear orientation is typically between 45° and perpendicular to the substrate (Supplementary Figure 5B). Nuclear orientations perpendicular to the basement membrane are even more pronounced in the *in vivo* intestine^77^. These differences in cellular force production and nuclear orientation likely influence the mode of mechanosensing and downstream responses in single cells vs. polarized epithelia, consistent with our findings. Interestingly, a recent study using human embryonic and pluripotent stem cells has demonstrated that mechanosensing at the level of global chromatin compaction can serve as a mechanical priming mechanism to modulate biochemically driven differentiation, similar to our observations^78^. Additional future studies will be required to determine whether mechanosensing in polarized intestinal epithelial cells also involves changes in chromatin organization.

Inhibition of STAT6 or YAP suppressed secretory cell hyperplasia in both IL-13- and stiffness-driven contexts (Figure 1G,H; Figure 5A,B), highlighting the essential role of both STAT6 and YAP in this process. YAP localization was predominantly cytoplasmic on soft substrates, became partially nuclear at intermediate stiffness, and reached high nuclear accumulation on stiff substrates. While nuclear elongation in response to high substrate stiffness could facilitate YAP nuclear accumulation via increased nuclear envelope tension and pore permeability as shown in previous studies^71,79^, our data indicate that YAP nuclear translocation is downstream of STAT6 (Figure 5C-F). This suggests that YAP nuclear accumulation in polarized epithelial tissues may not only be regulated directly by nuclear mechanics, but may involve more complex regulatory mechanisms. Our results indicate that STAT6 activation primes actomyosin contractility and cytoskeletal rearrangement by promoting MLCK1 expression, leading to increased MLC phosphorylation. This implies a positive feedback loop by which activation of STAT6—either via the upstream IL-13 receptor or via mechanosensing— increases actomyosin contractility, as evidenced by increased MLC phosphorylation and increased traction force generation (Figure 3A-C, Figure 6B,D), further potentiating the mechanosensing response. We predict that together, these changes promote mechanosensitive YAP activation, resulting in goblet cell hyperplasia as well as reduced barrier function due to competition between adhesion types (Figure 6E).

Our Myosin-2 inhibition experiments revealed that actomyosin contractility is not simply a downstream effect of substrate mechanosensing but is also functionally required for secretory cell differentiation driven by both IL-13 and high substrate stiffness (Figure 2C,D). Myosin-2 inhibition not only suppressed secretory fate acquisition but also impaired nuclear translocation downstream of both IL-13 and mechanosensing, indicating a requirement for actomyosin contractility in both signaling contexts (Figure 2E,F). Additionally, STAT6 inhibition reduced cortical myosin localization and secretory differentiation even on stiff substrates, indicating that activated STAT6 reinforces contractility (Figure 1G,H; Figure 2G,H). This reciprocal relationship implies a positive feedback loop between actomyosin tension and STAT6 signaling, where mechanical cues initiate cytoskeletal remodeling that enhances STAT6 activation, which in turn sustains the contractile machinery. Our biochemical analysis suggests that this feedback is mediated by STAT6-dependent expression of MLCK1. MLCK1 was previous identified as a STAT6 target and was implicated in compromising barrier function^66^. However, the mechanism was previously unclear. Our results provide a mechanism for this observation, namely that the resulting increase in actomyosin contractility leads to increased mechanical stress at cell-substrate adhesions and reduced tension at cell-cell adhesions. Extending this feedback model, YAP nuclear localization could be induced by either high stiffness or IL-13, but could be abolished by inhibition of either Myosin-2 or STAT6. This suggests that YAP is downstream of the feedback cycle between Myosin-2 and STAT6, and that direct binding of YAP and STAT6 could lead to a cooperative nuclear import mechanism. Together, these findings position actomyosin and STAT6 as central integrators of mechanical and inflammatory signals, linking substrate stiffness and IL-13 to coordinated activation of YAP in regulating epithelial stem cell differentiation. Our findings also suggest that mechanosensing mechanisms in polarized epithelial cells, which are currently not well understood, may differ from those previously identified in single cell in culture, warranting more investigation in the future. Taken together, the mechano-inflammatory framework identified here not only enhances our understanding of epithelial fate decisions but also opens new therapeutic avenues targeting contractility, STAT6, or YAP in chronic intestinal disease.

## Data availability

Primary data will be shared by the corresponding author upon reasonable request.

## Code availability

Custom code written for 3D segmentation as well as the StarDist segmentation model will be made available open-source via github upon publication (https://github.com/agclark12; https://github.com/Clark-Lab-Stuttgart). Other custom code for image and data analysis will be shared by the corresponding author upon reasonable request.

## Acknowledgments

We thank Markus Morrison and the members of the Clark lab for insightful discussions and critical reading of the manuscript. The authors gratefully acknowledge the Technology Platform “Cellular Analytics” of the Stuttgart Research Center Systems Biology for their support and assistance in this work. We also thank the staff of the animal facility of the Institute of Cell Biology and Immunology at the University of Stuttgart for their care of mouse stocks and support for experimental procedures. S.S.,

T.N. and A.G.C. were supported by the Federal Ministry of Education and Research (BMBF) and the Baden-Württemberg Ministry of Science (MWK-BW) as part of the Excellence Strategy of the German Federal and State Governments (NWG-GastroTumors to A.G.C.). C.M. and U.S.S. were supported by the cluster of excellence 3DMM2O (EXC 2082/1-390761711) funded by the Deutsche Forschungsgemeinschaft (DFG, German Research Foundation). This work was also supported by the Terra Incognita Program from the University of Stuttgart.

## Author contributions

S.S. and A.G.C. designed the research and wrote the manuscript. S.S. carried out the experiments and image analysis. A.G.C. wrote and tested custom software for image and data analysis. T.N. provided assistance for some experiments. C.M. and U.S. wrote and tested custom software for performing TFM analysis. M.T.-R. performed and analyzed some of the 3D organoid immunostaining experiments. K.K. generated ground truth segmentations for training the 3D nuclear segmentation model. S.E. designed and optimized protocols for microscopy. All authors discussed the results and manuscript.

## Methods

### Reagents

All critical reagents used in experimental procedures are listed in the supplementary information (Supplementary Table 1).

### Animals

To generate mouse intestinal organoids, C57BL/6J or C57BL6/N (male or female) mice ranging from eight weeks to six months old were used. All mice were housed under specific-pathogen-free conditions (12 h light/dark cycle, ad libitum access to food and water). Animals were euthanized by CO2 asphyxiation followed by cervical dislocation, and tissues were collected immediately post-mortem. mT/mG mice^80^ were purchased from JAX via Charles River Europe and imported in to the local mouse quarantine facility. mT/mG mice were housed for 1-2 weeks prior to sacrifice for the purpose of harvesting intestinal organs to generate organoids. All housing procedures and euthanasia were performed in accordance with local and regional laboratory animal regulations and experiments were approved by the local ethics board (relevant project IDs approved by the University of Stuttgart ethics board: IIG§4_2_2021.Clark, IZI§4-3-2022, IIG§4_1_2023, IZI§4_3_2024).

### Small intestinal crypt isolation

Isolation of small intestinal crypts was performed as in previous studies^31,41^. Immediately following sacrifice, the small intestine (SI) was removed by incising distal to the pylorus and proximal to the caecum. The mesentery was stripped, luminal contents flushed with ice-cold Ca²⁺/Mg²⁺-free PBS (PBS-0), and the SI was opened longitudinally. After washing twice with ice-cold PBS-0, 15 mins each at 4 °C, the tissue was cut into 2-3 mm fragments and incubated in ice-cold Dissociation Solution (2 mM EDTA, 1 mM DTT, 10 mM HEPES, pH 7.4) for 30 min on a rocking platform at 4 °C. Crypts were released by vigorous pipetting: the first two fractions (villus-enriched) were discarded, and the remaining fractions (crypt-enriched) were filtered (100 µm, then 70 µm) and pelleted by centrifugation (300g, 5 min, 4 °C).

### 3D organoid culture

Crypt pellets were resuspended 1:1 in Cultrex Basement Membrane Extract (BME) and Resuspension Buffer (4° DMEM/F12 + 2% Anti/Anti), and 50 µL domes were plated in 24-well plates. After polymerization at 37°C for 30 min, 350 µL of serum-free ENR medium (Advanced DMEM/F-12, 1x GlutaMAX, 10 mM HEPES, 1x Pen/Strep, 1x B27 minus vitamin A, 1x N-2, 50 ng mL^-1^ EGF, 100 ng mL^-1^ Noggin, 500 ng mL^-1^ R-spondin-1) was added. ENR Medium was refreshed every 2-3 days. Cultures were passaged every 4-5 days by mechanical dissociation and re-embedding in fresh BME (500xg, 5 min, 4°C).

### Preparation of organoid monolayers

Preparation of organoid monolayers was adapted from previous work^31^. Glass-bottom culture dishes (35 mm diameter) were pre-coated by incubating with 300 μl water and 150 μl silane APTMS for 15 minutes. After removing excess silane and washing twice with water, dishes were soaked in water for 10 minutes with agitation, then dried with compressed air. The dishes were subsequently incubated for 30 minutes with 0.5% glutaraldehyde in PBS. Following extensive washing and drying, dishes were stored dry at 4°C until use.

Polyacrylamide (PAA) gels were prepared on the pre-coated glass bottom dishes using established and validated formulas (Supplementary Table 2)^31,81^. Then, 16 μl of gel solution was applied onto glass dishes and gently overlaid with an 18 mm diameter round coverslip. The gels were allowed to polymerize for exactly 60 min at room temperature, after which the coverslips were removed in PBS using a razor blade and forceps. Gels were washed briefly with PBS and stored at 4°C in PBS for up to one week.

Polydimethylsiloxane (PDMS) stencils were fabricated by mixing the base and curing agent at a 10:1 ratio, degassed via centrifugation (3200 RPM, 2 min, RT), poured into plastic petri dishes to a height of 2mm, and cured at 80°C for at least 1 hour to overnight. The cured PDMS sheets were cut into ∼2 cm square shapes using a razor blade and perforated with three 1 mm holes using a biopsy punch. Stencils were stored dry and cleaned with 70% ethanol before reuse. PDMS stencils were passivated by incubation in 2% (w/v) Pluronic F-127 solution in PBS for 1 hour at room temperature with occasional swirling, followed by briefly rinsing with 70% ethanol and sterile water. Afterwards, stencils were washed twice with PBS, dried with compressed air, and air-dried for an additional 15 minutes.

One day before plating, PAA gels were activated by applying 50 μl of 2 mg/ml Sulfo-SANPAH solution and UV-irradiated (365 nm) using a UV Bench Lamp (Analytik Jena, Jena, Germany) for 7.5 minutes. Sulfo-SANPAH solution was then removed, and gels were washed with 10 mM HEPES 3 times for 2mins each followed by one time PBS wash for 2mins on an orbital shaker, then air-dried briefly. Activated gels were used within 30 minutes. Passivated PDMS stencils were placed on the activated gels. 10 μl of cold ECM coating solution containing collagen-I (250 μg/ml) and laminin-1 (100 μg/ml) in 10 mM HEPES was pipetted onto the stencil holes and incubated overnight at 4°C.

Organoids cultured in 3D were passaged one day prior to plating. On the plating day, BME domes containing organoids were incubated in Organoid Harvesting Solution at 4°C for 1 hour with gentle shaking. Organoids were then gently dissociated from the BME dome, centrifuged (500xg, 5 min), washed twice with PBS for one time (centrifuged at 500xg, 5min), and resuspended in ENR + Y-27632 medium (15 μl per gel). 15 μl of this suspension was seeded onto the ECM-coated regions of each PAA gel. After 1-2 hours incubation at 37°C and 5% CO2, 950 μl of fresh ENR medium was carefully added. Organoid monolayers were then used for imaging 2-4 days after plating.

### Cytokine and pharmacological inhibitor treatments

Organoid monolayers were treated with recombinant murine IL-13 (20 ng mL^-1^) for 48h (secretory differentiation, myosin/YAP localization, nuclear shape) or 24h (STAT6 localization). For inhibition assays, cultures were treated with AS1517499 (STAT6 inhibitor, 1 µM), verteporfin (YAP inhibitor, 2 µM) or Blebbistatin (myosin-II inhibitor, 50 µM) ± IL-13 for 48h. For Western Blot experiment to check for the expression of pSTAT6, the organoid monolayers were treated with IL-13 for 2h. For Western Blot experiment to check for the expression of YAP, the organoid monolayers were treated for 48h.

### Indirect immunofluorescence staining and confocal microscopy

Samples were fixed in 4% paraformaldehyde for 25 min at room temperature, permeabilized using 1% Triton X-100 for 5 min at room temperature, blocked in PBS + 5% BSA + 0.05% Tween-20 (PBS-T) for 1 h at room temperature, and incubated with primary antibodies (1:100 in PBS-T, overnight, 4°C; see Table S1) followed by AlexaFluor-conjugated secondary antibodies (1:1000, overnight, 4°C) and DAPI (0.5 µg mL^-1^, 30 min). Phalloidin-AlexaFluor647 (1:2000) was added with secondary antibodies. Samples were washed thoroughly three times with PBS-T (0.1% Tween-20) between steps. After washing, one drop of Fluoromount-G was added on the top of the sample and mounted with a coverslip.

All immunofluorescence samples were imaged using an LSM980 Airyscan 2 microscope (Carl-Zeiss Microscopy, Germany) equipped with a LD LCI Plan-Apochromat 40x/1.2 Water immersion objective. Images were acquired in 4Y multiplex mode using the following diode laser/beamsplitter (detection range) combinations: DAPI channel: 405 nm/300-720nm, green channel: 488 nm/495-550 nm, red channel: 561 nm/573-627 nm, Phalloidin channel: 640 nm/574-720 nm. Airyscan raw images were 3D-processed using Zen Blue software (v3.3).

### Western blotting

3D organoids were removed from BME domes as described above for plating organoid monolayers including centrifugation at 500x g for 5 min. For organoid monolayers, the monolayer was removed from the PAA gel by flushing with ice-cold PBS and centrifuged at 500xg for 5 min. Following centrifugation for 3D organoids or organoid monolayers, RIPA buffer supplemented with an EDTA-free protease inhibitor cocktail was added to the pellet to lyse the organoids. The lysate solution was kept on ice for 15 mins before centrifuging at 16000x g for 15 mins at 4°C. The supernatant was removed and protein concentration was determined by Bradford assay. 20 µg protein per lane was resolved by SDS-PAGE, transferred to a nitrocellulose membrane, blocked (5% Roti Block in TBS-T), and probed with primary antibodies overnight at 4C followed by HRP-conjugated secondary antibodies for 2 hours at room temperature. Bands were visualized using ECL substrate, imaged using a Fusion Solo documentation station (Vilber) and quantified by densiometry using Fiji^82^. Raw uncropped images of the membranes can be found in the supplementary information (Supplementary Figure 7).

### Co-immunoprecipitation

3D organoids were cultured for 48 h prior to IL-13 treatment. Organoids were harvested as described for Western blotting and lysed in reaction tubes with 200-250 μl IP lysis buffer (150 mM NaCl, 20 mM TRIS-HCl pH 7.5, 1 mM EDTA, 1 mM EGTA, 1% (v/v) NP-40, 1x cOmplete protease inhibitor cocktail, 1x PhosSTOP phosphatase inhibitor; pH 7.6) for 30 min on ice. The reaction tubes were then centrifuged at 16000x g and 4°C for 15 min. Protein concentration was determined via Bradford assay as for Western blot.

Protein G agarose beads were prepared by centrifuging 200-250 μl of bead solution at 2500x g and 4°C for 2 min to obtain at least 20 μl of pure beads. The supernatant was then discarded, and the beads were washed by adding 1 ml Tris-HCl (pH 7.4). After re-centrifugation (2500x g, 4°C, 2 min) and discarding the supernatant, 1 ml of IP buffer (150 mM NaCl, 20 mM TRIS-HCl pH 7.5, 1 mM EDTA, 1 mM EGTA, 1% (v/v)

NP-40; pH 7.6) was added to the beads. The solution was then centrifuged again (2500x g, 4°C, 2 min). The supernatant was removed, and a volume of IP buffer equal to the volume of the remaining beads was added to the reaction tube. Beads were stored for up to two weeks at 4°C.

Full lysate samples (input) were prepared as described in the Western Blot protocol (20 μg protein per sample). For immunoprecipitation, 500-1000 μg of protein was used, and the lysate was filled to a total volume of 1 ml with IP lysis buffer before adding 2 μg anti-Stat6 antibody and mixing on a rolling shaker for 2 h at 4°C. Meanwhile, the previously prepared bead solution was equally distributed to new reaction tubes with 20 μl of beads per tube. After shaking, the lysates with anti-Stat6 antibody were transferred to the prepared beads, carefully inverted, and mixed on a rolling shaker at 4°C for 1.5 h. The samples were then centrifuged at 2500x g and 4°C for 2 min. The supernatant was discarded, and the beads were washed 3x with IP lysis buffer (800 μl per reaction tube) to remove non-specifically bound proteins. The beads were then resuspended in 20 μl 2x Laemmli buffer with DTT and incubated on a heat block at 95°C for 10 min to denature the proteins, and then frozen at -20°C or loaded directly onto a SDS gel. Western blotting for YAP was performed as described above.

### Laser ablation assay

Monolayers derived from mT/mG reporter mice were imaged on an inverted Stellaris8-DIVE confocal LSM (Leica microsystems, Germany), equipped with a Whitelight laser (range 440nm-790 nm) and an Insight X3 dual Ti-sapphire laser (Spectraphysics, CA, USA) using a HC PL IRAPO 40x/1,1 W objective. mT/mG channel (excitation: 550 nm, emission 562-732 nm) was acquired with a frame rate of 3 s^-1^ for 35 cycles. After 5 cycles, individual cell-cell junctions were targeted with the 1045 nm fixed line of the Ti-Sapphire laser at one bleaching point for 10 ms at 25% transmission. The time series were then analyzed in Fiji.

### Quantification of Myosin Localization in Goblet Cells

The subcellular distribution of pMLC was quantified specifically in goblet cells. To assess the basal-to-apical distribution of myosin signal intensity, regions of interest (ROIs) around individual goblet cells were manually selected based on phalloidin morphology. Z-axis intensity profiles were generated by plotting mean fluorescence intensity values across the entire apical-basal axis of each cell. Cortical myosin enrichment was measured using linescan analysis across the apical cross-section of goblet cells. A standardized line width of 30 pixels (approximately 2.5 μm) was drawn through the diameter of the apical goblet cell region, perpendicular to the cell’s apico-basal axis. Mean fluorescence intensity was measured along this line to generate intensity profiles revealing both cortical and cytosolic signal distribution. Cortical signal intensity was defined as the average of the two peak fluorescence values detected at the lateral edges of the line scan, corresponding to the cell cortex regions. Cytosolic signal intensity was calculated as the average of all intermediate values between these cortical peaks, representing the intracellular compartment. The cortical:cytosolic ratio was subsequently calculated by dividing the cortical signal intensity by the cytosolic signal intensity for each analyzed cell (Supplementary Figure 3C).

### Quantification of UEA I+ Secretory Cells

Identification and quantification of secretory cell frequencies was determined based on DAPI and UEA I staining. First, nuclear segmentation was performed using the StarDist framework for python^83,84^. To train a custom 3D segmentation model in StarDist, we created a ground-truth data set by manual segmentation. To achieve this, DAPI-stained 3D organoids and organoid monolayers were imaged with the same parameters as described above for immunofluorescence. Each of the ∼30 images was square with 300 px side length and 36 z-planes and contained ∼50-200 nuclei. The relative size of each nucleus was ∼30 px in diameter. Each nucleus was manually segmented using the LabKit plugin^85^ in Fiji. This annotation was performed in a lazy manner, whereby the perimeter of the nucleus in ∼5 selected z-planes (including the bottom and top frames where the nucleus was visible) was annotated. Using a custom script in python, contours were filled in the annotated z-planes. We next used ITK-SNAP^86^ to perform an interpolation of the segmented regions to fill in the non-annotated z-planes, which resulted in the full annotation of each nucleus in 3D. The resulting ground-truth data set was used to train the StarDist segmentation model, including using image augmentation (rigid image rotation, intensity modification) to increase the number of images used for training. Training was performed using the TensorFlow machine learning platform^87^ and CUDA^88^. For training, 20% of the images were selected at random to serve as a validation set, and training was performed to 300 epochs. Loss over these epochs was observed to decrease exponentially and plateau by live tracking using TensorBoard^87^.

The resulting segmentation model was then used to segment nuclei in 3D image stacks of 3D organoids and organoid monolayers using similar imaging conditions as for the training data set. To generate perinuclear regions to determine cytoplasmic intensities, nuclear segmentation masks were expanded using a 3D spherical structuring element (3 px diameter) over 3 iterations and subtracting the segmented nuclear regions. To determine whether each cell contained mucus granules (characteristics of secretory goblet cells), the mean UEA I intensity was determined in the perinuclear shell for each cell using the MorphoLibJ plugin^89^ in Fiji. For each experimental sample, fluorescence intensities across all cells were combined into histograms, which were then used to identify the bimodal distribution characteristic of mixed epithelial cell populations containing UEA I negative (absorptive enterocytes) and UEA I positive (goblet cells) populations. This enabled accurate sample-specific thresholding to account for differences in staining intensities across samples. Cells with mean fluorescence intensities above this threshold were classified as UEA I positive goblet cells, and the frequency of goblet cells was determined by taking the ratio of UEA I goblet cells to the total cell number from nuclear segmentation (Supplementary Figure 1).

### Quantification of STAT6 and YAP Nuclear Translocation

Automated nuclear and perinuclear cytoplasm segmentation was performed using StarDist for DAPI-stained nuclei as described for the quantification of secretory cells. From the nuclear and perinuclear cytoplasm regions, mean intensities of STAT6 or YAP were determined using MorphoLibJ in Fiji. For each cell, the nuclear: cytoplasmic ratio was defined as the mean nuclear intensity divided by the mean perinuclear cytoplasmic intensity.

### Quantification of Nuclear Morphology

Automated nuclear segmentation was performed using the StarDist for DAPI-stained nuclei as described above. Image preprocessing was performed prior to segmentation to optimize detection accuracy. DAPI-stained z-stack images were acquired with consistent imaging parameters (typically 0.8 μm z-step intervals) to ensure adequate sampling of nuclear volume. Image stacks were deconvolved using appropriate point spread function parameters to reduce out-of-focus blur and enhance edge definition. Noise reduction was applied using Gaussian filtering (σ = 0.5-1.0 pixels) when necessary, while preserving edge information critical for accurate boundary detection. After the nuclei were segmented, the nuclear morphometrics were extracted using MorphoLibJ in Fiji.

### Traction Force Microscopy

For TFM experiments, PAA gels of different stiffness were polymerized as described above, but with the addition of fluorescent beads (Supplementary Table 2). Organoid monolayers were cultured on PAA gels containing beads as for standard PAA gels described above. To determine bead displacements, the organoid monolayers and fluorescent bead-containing gels were imaged using confocal microscopy. Images were acquired on a AxioObserver SD Spinning Disk microscope (Carl-Zeiss microscopy, Germany) equipped with a LD LCI Plan-Apochromat 25x/0.8 Oil objective in brightfield and red fluorescence channels (excitation: 561 nm, emission filter 600/50 nm). Following the acquisition of one image stack, the cells were detached by treating with 0.25% trypsin for 5mins. After ensuring that all cells were successfully removed, a second image stack was acquired. Bead displacements were interpolated onto a regular grid from features tracked by a Kanade-LucasTomasi (KLT) optical flow algorithm, and tracked features were selected based on fitness^90^. A background set of tracked features was additionally used, wherein the fittest features of regularly spaced windows were tracked. This background prevented extrapolation in regions with lower feature fitness and increased the accuracy of calculated displacement fields. To correct for substrate drift between the moments of image acquisition, the average of calculated displacements was subtracted from the final displacement field. Traction maps were reconstructed from the calculated displacement fields via the inverse Green’s Function method, more widely known as Fourier Transform Traction Cytometry (FTTC)^91–93^. To stabilize the inverse and ill-posed problem of TFM, 0^th^ order Tikhonov regularization was used to reconstruct traction forces^94^. The regularization parameter for this procedure was estimated via a Generalised Cross Validation (GCV) technique^92^. A Tukey filter with small taper was applied to the displacement field to avoid force monopoles at the edges of the image, where cells may only be partially inside the frame.

To generate average radial linescans from the center of crypts from resulting traction stress fields, binary masks were generated manually from the pre-trypsin monolayers using Fiji. Using custom scripts in python (Python Software Foundation, https://www.python.org/), tangential and radial stresses were extracted by cubic interpolation of the stress fields along linescans at each of 360° angles originating from the crypt center toward the edge of the crypt and extending to 1.5 times the crypt radius. The radial linescans were then averaged by taking a mean along the linescan length to produce an average linescan for each crypt. To average linescans across crypts, the spatial coordinates of the mean crypt linescans were normalized such that the center of the crypt was at position x=0, and the edge of the crypt was at position x=1. The mean linescans were then averaged by taking the mean along this rescaled spatial coordinate to produce average linescans for each experimental condition. Custom python scripts used for the determination of linescan coordinates, interpolation and linescan averaging used the following additional packages: matplotlib^95^, numpy ^96^, pandas^97^, scikit-image^98^, scipy^99^ and tifffile^100^.

### Lucifer Yellow organoid permeability assay

The organoid-based epithelial permeability assay is based on a previously published protocol^61^. 3D Organoids were passaged as described above, and 25 µL of the organoid BME mixture was dispensed into each compartment of a 4-chamber glass-bottom dish. After allowing the organoids to settle and adhere for 24 hours, the culture medium was replaced with treatment medium as described above. Following treatment, organoids were transferred to the microscope stage and maintained at 37°C with 100% relative humidity and 5% CO2 throughout imaging. Live imaging was performed using a Cell Observer Z1 epifluorescence microscope (Carl Zeiss Microscopy, Germany) equipped with an Axiocam 503 camera and an EC Plan-Neofluar 20x/0.5 objective. Lucifer Yellow fluorescence intensity was acquired using a 470nm LED light source in combination with a Zeiss filterset 62. Just prior to imaging imaging, Lucifer Yellow was added to the culture medium to a final concentration of 1 mM. Timelapse imaging was performed every 7.5 minutes over a 5-hour period to monitor dye permeability into the organoid lumen. Organoid positions were tracked during the experiment using an automated stage with multi-position imaging. For image analysis, Fiji was used to extract the mean fluorescence intensity of the organoid lumen and background. Relative luminal fluorescence (RLF) was calculated as the product of the relative lumen fluorescence at each timepoint and its initial value, divided by the product of the background fluorescence and its initial value.

### Particle Image Velocimetry Analysis (PIV)

The migration of cells within the crypt-like compartment was quantified using particle image velocimetry (PIV) analysis. PIV computations were performed using python-based software developed in the working group^101^, based on openpiv^102^. Images were obtained using an AxioObserver SD Spinning Disk microscope in the brightfield channel as described above for TFM experiments with time intervals of 10 min for 24h.

### Data Visualization, Statistical Analysis and Image Analysis

Generation of plots and statistical analysis was performed using GraphPad Prism and matplotlib^95^. For boxplots, dots represent individual measurements as described in the figure legends. Boxes represent the 25^th^ to 75^th^ percentiles of the data. The line inside of the box is the median. Whiskers extend to the minimum and maximum value. The Large Language Model ChatGPT was used to assist in writing custom python functions for image analysis. All code segments generated by ChatGPT were tested and validated.

