## Supplementary Information for "Mechanosensing and IL-13 Signaling Synergistically Modulate Intestinal Stem Cell Differentiation via STAT6 and YAP"

### Supplementary Figures

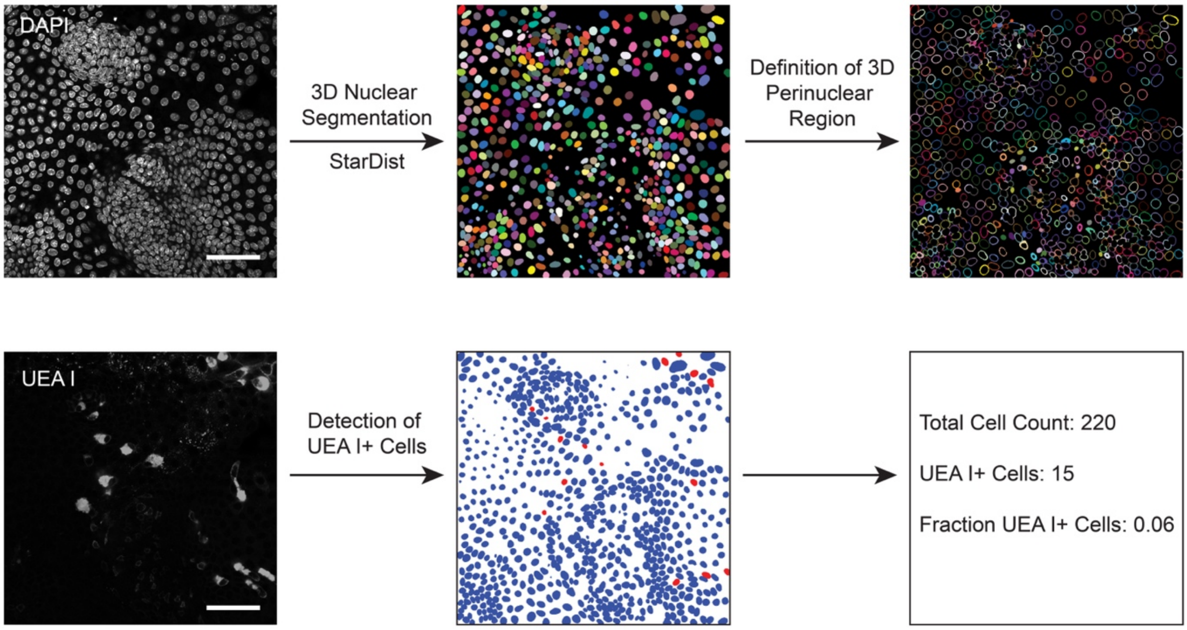


**Supplementary Figure 1: Description of image analysis workflow for quantification of UEA I+ cells.** Nuclei from 3D imaging stacks from DAPI-stained organoid monolayers or 3D organoids are segmented in 3D using StarDist based on a segmentation model from a custom ground-truth data set. The perinuclear cytoplasm region of each cell is then defined by expanding each nuclear label and removing the nuclear volume. Using the UEA I staining micrographs, the UEA I fluorescence intensity is determined for each perinuclear region, and the mean perinuclear intensity is thresholded in a semi-automated manner across all cells to determine the fraction of UEA I+ cells. SB: 50µm.

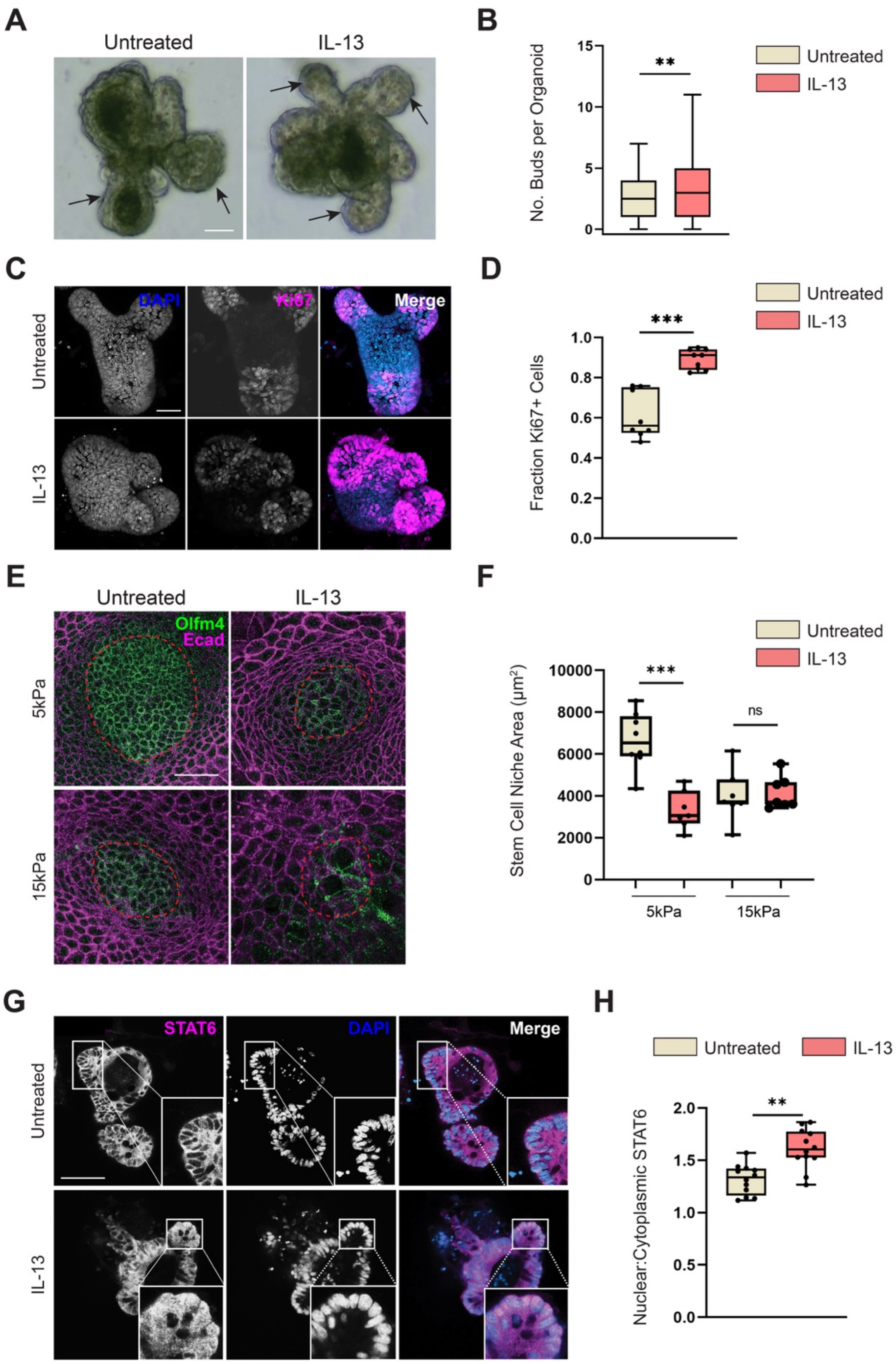


**Supplementary Figure 2: Substrate Stiffness and IL-13 affect intestinal stem cell organization and differentiation. A.** Brightfield micrographs of 3D organoids with or without IL-13 treatment. Black arrows indicate in-plane buds. SB:20μm. **B.** Quantification of number of buds per organoid for conditions in *A*. Each dot represents one organoid from *n* = 154, 149 organoids and *N* = 4, 4 independent experiments. **C.** Representative micrographs of 3D organoids treated with or without IL-13 and immunostained with Ki67 to identify actively dividing cells and stained with DAPI to label cell nuclei. Images are maximum intensity projections of z-stacks. SB: 100μm. **D.** Quantification of Ki67+ cell fraction for conditions in *C*. Each dot represents one organoid from *n* = 8, 9 organoids and *N* = 2, 2 independent experiments. **E.** Representative micrographs of organoid monolayers cultured on intermediate (5 kPa) or stiff (15 kPa) polyacrylamide (PAA) gels, treated with or without IL-13 and immunostained for Olfm4 and E-cadherin (Ecad). Images are max projections of z-stacks. Red dashed lines indicate the crypt-like compartments. SB: 20μm. **F.** Quantification of Olfm4+ stem cell niche area for conditions in *E*. Each dot represents one organoid monolayer from *n* =, 8, 7, 7, 7 organoid monolayers and *N* = 2, 2 independent experiments. **G.** Representative micrographs of 3D organoids treated with or without IL-13 and immunostained with STAT6 to identify actively dividing cells and stained with DAPI to label cell nuclei. Images are maximum intensity projections of z-stacks. SB: 100μm. **H.** Quantification of nuclear:cytoplasm intensity ratio of STAT6 for conditions in *E*. Each dot represents one organoid from *n* = 12, 12 organoids and *N* = 2, 2 independent experiments. For panel *F*, one-way ANOVA, $p<0.001$. Pairwise tests performed using Welch's t-test (*B,D,H*) or Tukey's HSD post-hoc test (*F*; **$p<0.01$, ***$p<0.001$, ns:$p>0.05$).

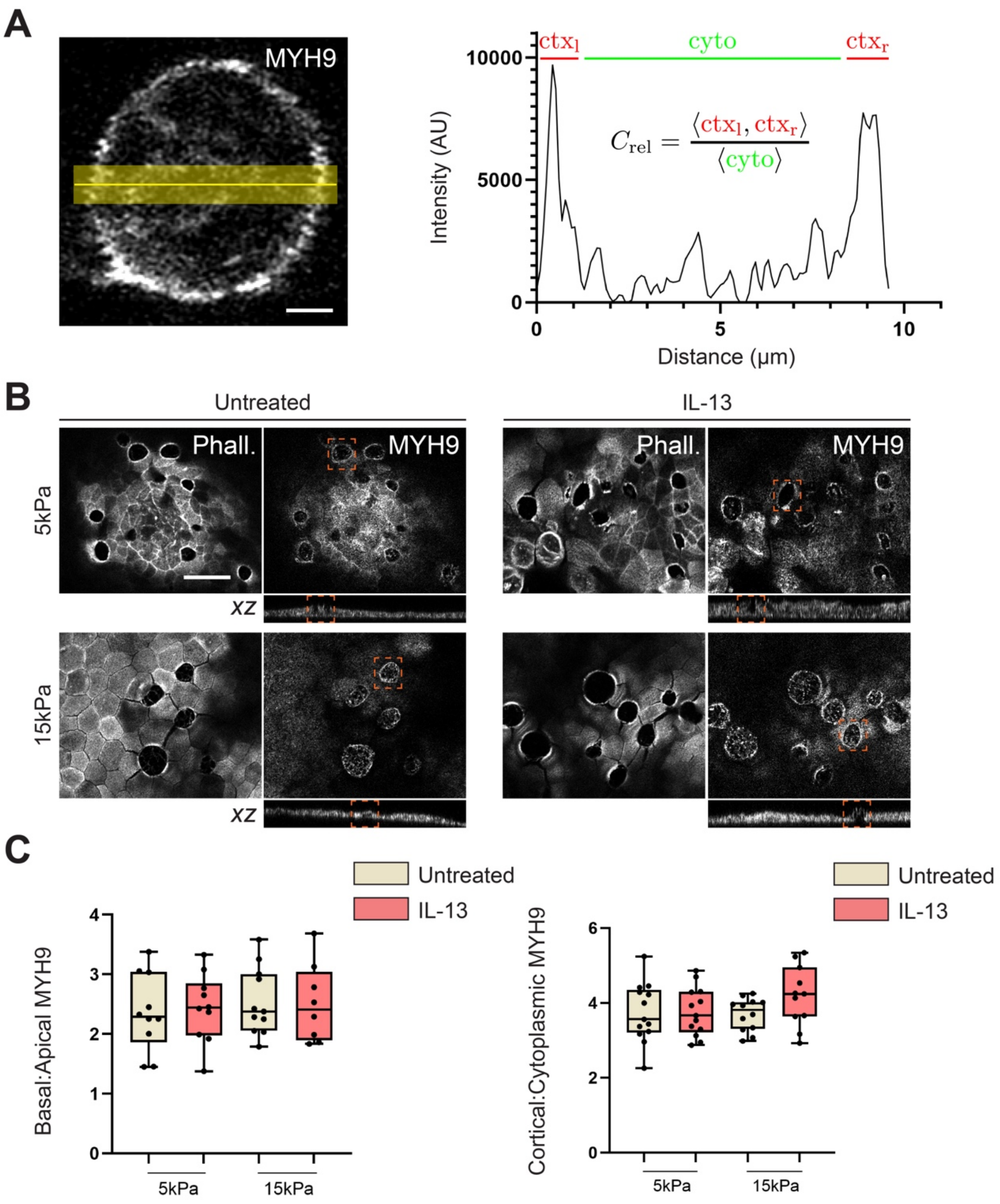


**Supplementary Figure 3: MYH9 localisation is altered in response to IL-13 and substrate stiffness A.** Linescan analysis workflow for measuring cortex to cytoplasm ratio for localization of Non-muscle myosin heavy chain 2A (MYH9) or phosphorylated myosin-2 regulatory light chain (pMLC; Figure 2A). SB:10μm. **B.** Representative micrographs of organoid monolayers cultured on intermediate (5 kPa) or stiff (15 kPa) polyacrylamide (PAA) gels, treated with or without IL-13 and immunostained for MYH9 and stained for filamentous actin (F-actin) using phalloidin. The x-z view shows the basal:apical MYH9 localisation ratio. SB: 20μm. **C.** Quantification of MYH9 intensity ratio of the cortical:cytoplasmic and apical:basal poles for conditions as in *A*. SB:5μm.

For Cortical:Cytoplasmic MYH9, each dot represents one goblet cell from *n* = 13, 13, 12, 11 goblet cells from 10, 10, 11, 8 organoid monolayers and *N* = 2, 2 independent experiments. For Basal:Apical MYH9, each dot represents one organoid monolayer from *n* = 10, 10, 11, 8 organoid monolayers and *N* = 2, 2 independent experiments. For panel *C*, one-way ANOVA was performed (p = 0.9151, 0.2882).

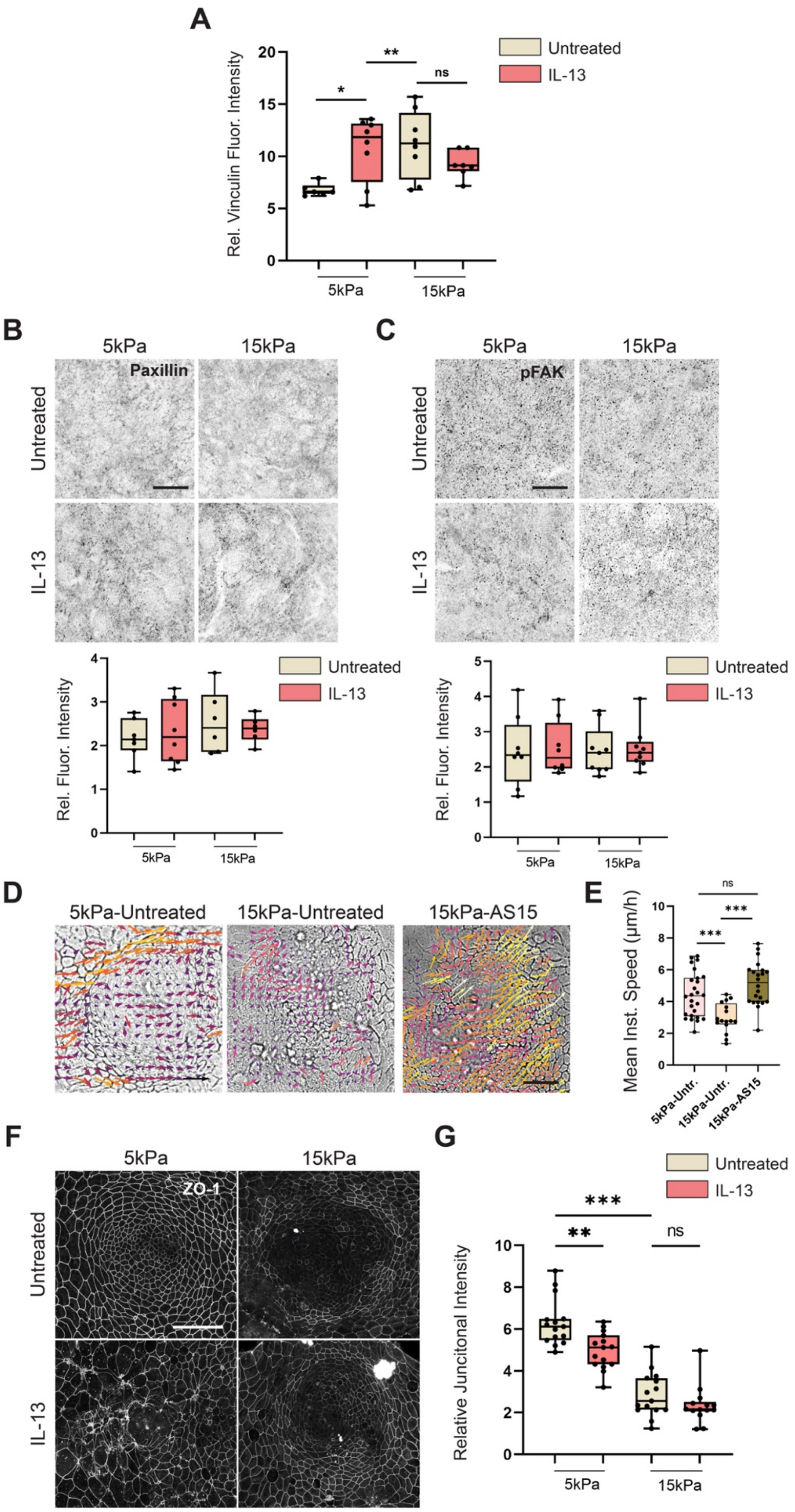


**Supplementary Figure 4. Substrate stiffness modulates cell-substrate and cell-cell adhesions. A.** Quantification of average relative fluorescence intensity of vinculin at focal adhesions (FAs) with respect to cytoplasmic background. Each dot represents one organoid monolayer from *n* = 6, 8, 8, 7 organoid monolayers and *N* = 3, 3, 3 independent experiments. **B-C.** Representative micrographs of organoid monolayers cultured on intermediate (5 kPa) or stiff (15 kPa) polyacrylamide (PAA) gels, treated with or without IL-13 and immunostained for FA components Paxillin (*B*) or phosphorylated Focal Adhesion Kinase (pFAK; *C*). *Below:* quantification of average relative fluorescence intensity of Paxillin/pFAK-enriched FAs with respect to cytoplasmic background. Images are max projections of z-stacks. SB: 20µm. For Paxillin fluorescence intensity (*B*), each dot represents one organoid monolayer from *n* = 7, 8, 6, 6 organoid monolayers and *N* = 2, 2 independent experiments. For pFAK fluorescence intensity (*C*), each dot represents one organoid monolayer from *n* = 8, 8, 9, 9 organoid monolayers and *N* = 2, 2 independent experiments. **D.** Representative micrographs from live imaging of organoid monolayers cultured on intermediate/stiff PAA gels, treated with or without the STAT6 inhibitor AS1517499 (AS15). Scale vector: 1µm/min. Scale Bar: 20µm. **E.** Quantification of mean instantaneous speeds extracted from timelapse imaging data and quantified by PIV for conditions in *C*. Each dot represents one crypt from *n* = 30, 16, 22 crypts and *N* = 2, 2 independent experiments. **F.** Representative micrographs of organoid monolayers cultured on intermediate/stiff PAA gels, treated with or without IL-13 and immunostained for the tight junction adaptor protein Zonula Occludens 1 (ZO-1). SB: 20µm. **G.** Quantification of ZO-1 intensity at cell-cell junction relative to cytoplasmic background from individual cell segmentation for conditions in *F*. Relative intensity was averaged across all junctions per image. Each dot represents one organoid monolayer from *n* = 15, 15, 15, 15 organoid monolayers and *N* = 2, 2, 2 ,2 independent experiments. For panels *A, B, C, E, G*, One-way ANOVA was performed (p = 0.0150, 0.7282, 0.9940, 2.0e-5, 8.6e-16). Pairwise tests performed using Tukey's HSD post-hoc text (*p<0.05, **p<0.01, ***p<0.001, ns:p>0.05).

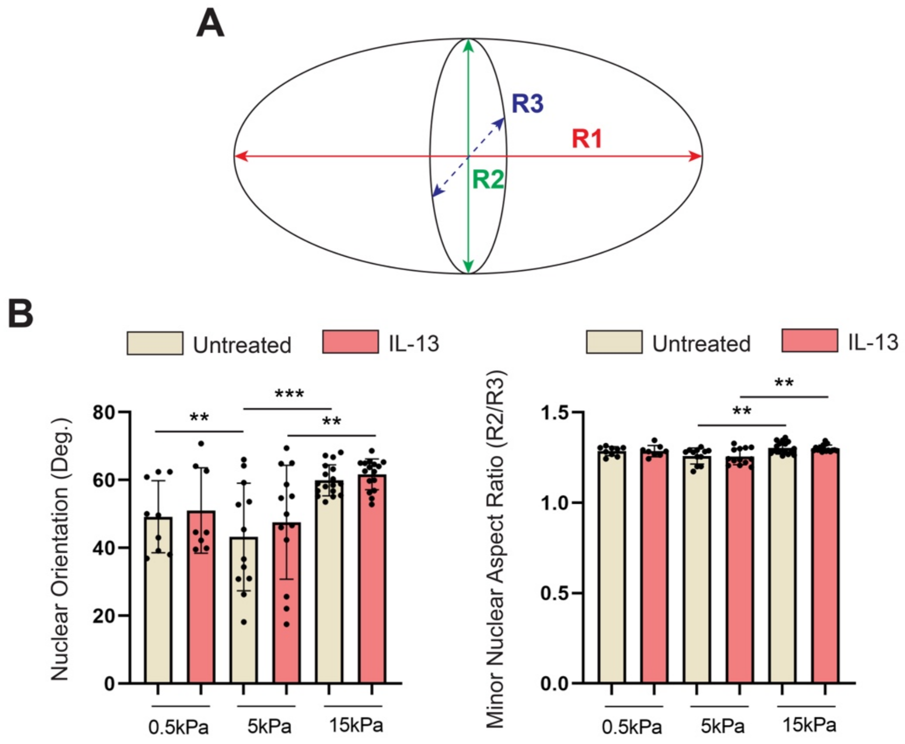


**Supplementary Figure 5: Quantification of nuclear morphometrics. A.** Schematic representation of the axes for ellipsoid fits of 3D segmented nuclei, indicating the radii along the three principal axes (R1, R2, R3). **B.** Quantification of nuclear orientation relative to the substrate (0º: parallel to the substrate, 90º: perpendicular to the substrate) and minor aspect ratio, R2/R3 (ratio of radii from two shortest principal axes) for cells in crypt-like regions for conditions in Figure 4E. For panel *B,* one-way ANOVA was performed (p = 0.0001, 0.0004). Pairwise tests performed using Tukey's HSD post-hoc text (**p<0.01, ***p<0.001, ns:p>0.05).

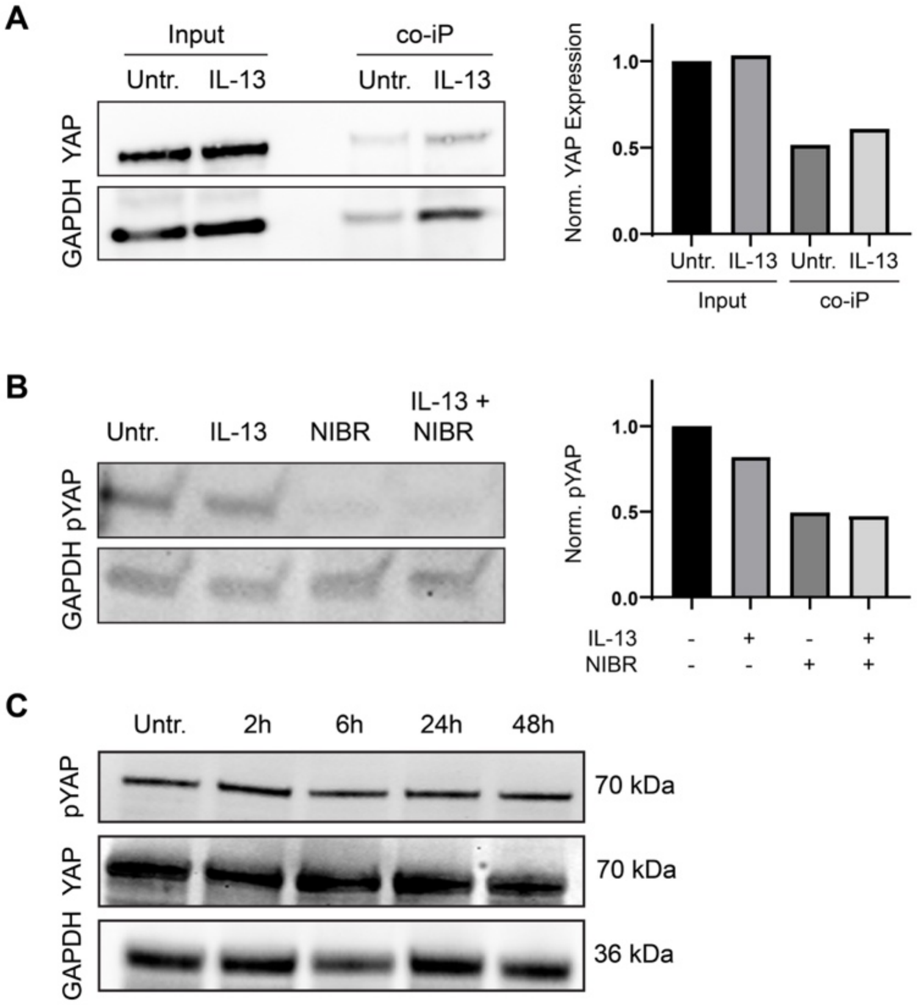


**Supplementary Figure 6: Interactions between STAT6, YAP and Hippo pathway. A.** Western blot analysis of YAP from whole 3D organoid lysates (input) or following immunoprecipitation of STAT6 (co-IP) with or without 24h IL-13 treatment. *Right*, quantification of Western blot by densiometry. **B.** Western blot analysis of pYAP levels following treatment with IL-13 and/or NIBR-LTSi, an inhibitor of the upstream Hippo kinase large tumor suppressor (LATS). *Right*, quantification of Western blot by densiometry. **C.** Western blot analysis of total and phosphorylated YAP levels at various time points following IL-13 treatment in 3D organoid monolayers.

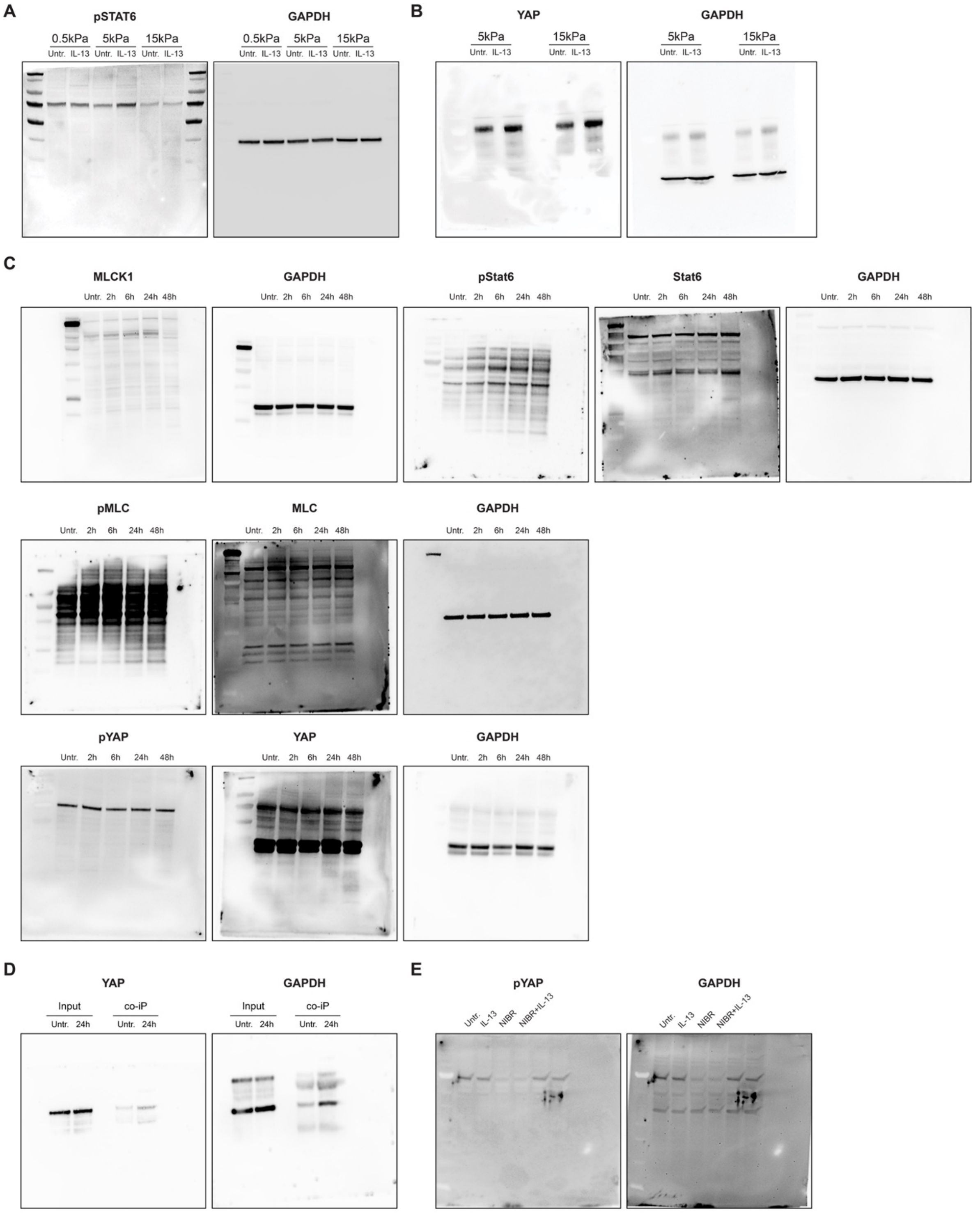


**Supplementary Figure 7: Uncropped Western blot images used in this study.** **A.** Uncropped Western blot for pSTAT6 used in Figure *1C*. **B.** Uncropped Western blot for YAP used in Figure *4C*. **C.** Uncropped Western blots used for time-course after IL-13 treatment used in Figure *6A-C* and Supplementary Figure *6C*. **D.** Uncropped Western blot from co-immunoprecipitation (STAT6 as bait, blotted for YAP) used in Supplementary Figure *6A*. **E.** Uncropped Western blot for pYAP following treatment with IL-13 and/or NIBR-LTSi, an inhibitor of the upstream Hippo kinase large tumor suppressor (LATS) used in Supplementary Figure *6B*.

**Supplementary Table 1. Critical reagents used in experimental procedures.**

| Reagent | Source | Product Number |
|---|---|---|
| **Antibodies/Fluorescent Dyes** | | |
| Rabbit-α-Olfm4 | Cell Signaling Technologies | 39141 |
| Rabbit-α-MLC2 | Cell Signaling Technologies | 8505 |
| Mouse-α-pMLC2 | Cell Signaling Technologies | 3675 |
| Rabbit-α-MYH9 | BioLegend | 909801 |
| Rabbit-α-MLCK1 | Abcam | ab76092 |
| Mouse-α-Occludin | Thermo Fisher Scientific | 740006M |
| Mouse-α-ZO-1 | Thermo Fisher Scientific | 33-9100 |
| Mouse-α-Paxillin | BD Biosciences | 610052 |
| Mouse-α-pFAK | Santa Cruz Biotechnology | sc-374668 |
| Mouse-α-Vinculin | Merck, Sigma-Aldrich | V9131 |
| Rabbit-α-YAP | Cell Signaling Technologies | 14074 |
| Rabbit-α-pYAP | Cell Signaling Technologies | 13008 |
| Rabbit-α-STAT6 | Abcam | ab32520 |
| Rabbit-α-pSTAT6 | Cell Signaling Technologies | 9361 |
| Rabbit-α-GAPDH | Cell Signaling Technologies | 2118 |
| Goat-α-Rabbit-IgG-AlexaFluor488 | Thermo Fisher Scientific | A-11008 |
| Goat-α-Mouse-IgG-AlexaFluor488 | Thermo Fisher Scientific | A-11001 |
| Goat-α-Rabbit-IgG-AlexaFluor568 | Thermo Fisher Scientific | A-11011 |
| Goat-α-Mouse-IgG-AlexaFluor568 | Thermo Fisher Scientific | A-11004 |
| Goat-α-Rabbit-IgG-AlexaFluor647 | Thermo Fisher Scientific | A-21244 |
| Goat-α-Mouse-IgG-AlexaFluor647 | Thermo Fisher Scientific | A-21235 |
| Goat-α-Rabbit-IgG-HRP | Merck, Sigma-Aldrich/Chemicon | AP307P |
| Goat-α-Mouse-IgG-HRP | Merck, Sigma-Aldrich/Chemicon | AP308P |
| Ulex Europaeus Agglutinin I (UEA I), Rhodamine conjugated | BIOZOL | VEC-RL-1062 |
| 4,6-diamidino-2-phenylindole (DAPI) | Thermo Fisher Scientific | D1306 |
| Phalloidin AlexaFluor647 | Thermo Fisher Scientific | A22287 |
| Lucifer Yellow | Thermo Fisher Scientific | L453 |
| Chemicals, peptides, and recombinant proteins | | |
| Recombinant Mouse Interleukin-13 (rmIL-13) | ImmunoTools | 12340133 |
| $PBS^{-/-}$ (1x DPBS without $Ca^{+2}/Mg^{+2}$) | Thermo Fisher Scientific, Gibco | 14190 |
| $PBS^{+/+}$ (1x DPBS with $Ca^{+2}/Mg^{+2}$) | Thermo Fisher Scientific, Gibco | 14040 |
| DMEM/F-12 | Thermo Fisher Scientific, Gibco | 10565018 |

| | | |
|---|---|---|
| Cultrex BME | Bio-Techne, R&D systems | 3533-010-02 |
| Culturex Organoid Harvesting Solution | Bio-Techne, R&D systems | 3700-100-01 |
| Fetal Calf Serum (FCS) | PAN-Biotec | P30-3031 |
| Gibco Antibiotika-Antimykotikum (100X) | Thermo Fisher Scientific, Gibco | 15240096 |
| GlutaMAX-1 (100X) | Thermo Fisher Scientific, Gibco | 35050061 |
| Murine EGF | Thermo Fisher Scientific, PeproTech | 315-09 |
| Murine FGF | Thermo Fisher Scientific, PeproTech | 450-33A |
| N2 Supplement (100X) | Thermo Fisher Scientific, Gibco | 17502048 |
| B27 Supplement (50X) | Thermo Fisher Scientific, Gibco | 17504044 |
| Y-27632 dihydrochloride | TebuBio | 282T1725 |
| AS1517499 | MedChemExpress | HY-100614 |
| (±)-Blebbistatin | Bio-Techne, Tocris | 1760 |
| Verteporfin | TebuBio | 282T3112 |
| NIBR-LTSi | MedChemExpress | HY-160769 |
| Ciprofloxacin | Merck, Sigma-Aldrich | 17850-5G-F |
| Metronidazol | Merck, Sigma-Aldrich | M1547 |
| DMSO | Carl Roth | 4720.4 |
| Trypsin/EDTA 0,5% 10x | Thermo Fisher Scientific, Gibco | 15400054 |
| 40% Acrylamide Solution | Bio-Rad | 1610144 |
| 2% Bis-Solution | Bio-Rad | 1610142 |
| N,N,N',N'-Tetramethylethylendiamine (TEMED) | Merck, Sigma-Aldrich | 1107320100 |
| FluoSpheres (0.2 µm, yellow-green) | Thermo Fisher Scientific, Invitrogen | 10513463 |
| 16% Formaldehyde (w/v) | Thermo Fisher Scientific | 28908 |
| Glutaraldehyde | Merck, Sigma-Aldrich | G6257 |
| (3-Aminopropyl) trimethoxysilane | Merck, Sigma-Aldrich | 440140 |
| Laminin 1 (Mouse) | Thermo Fisher Scientific | 23017015 |
| Rat Tail Collagen Type 1 | Corning | 354236 |
| HEPES (1M) | Thermo Fisher Scientific, Gibco | H3537 |
| Pluronic F-127 | Merck, Sigma-Aldrich | P2443 |
| Sulfo-SANPAH (Sulfosuccinimidyl-6- (4'-azido-2'-nitrophenylamino) hexanoat) | Merck, Sigma-Aldrich | 803332 |
| Triton X-100 | Merck, Sigma-Aldrich | T8787 |
| Tween-20 | Carl Roth | 9127.8 |
| Protein G Agarose | Thermo Fisher Scientific, Pierce | 20398 |

| | | |
|---|---|---|
| cOmplete protease inhibitor cocktail | Merck, Roche | 11697498001 |
| PhosSTOP phosphatase inhibitor | Merck, Roche | 4906845001 |
| ECL Western Blotting Substrate | Thermo Fisher Scientific, Pierce | 32109 |

**Supplementary Table 2. Composition of PAA Substrates.** *Note: fluorescent beads (FluoSpheres) were added for traction force microscopy experiments only. For other experiments, no beads were added, the volume of PBS was increased accordingly to ensure the same final volume.

| **Stiffness (kPa)** | **1x PBS (µl)** | **40% Acrylamide (µl)** | **2% Bis-acrylamide (µl)** | **Fluorescent Beads (µl)** | **10% APS (µl)** | **TEMED (µl)** |
|---|---|---|---|---|---|---|
| 0.5 | 434.75 | 50 | 7.5 | 5 | 2.5 | 0.25 |
| 5 | 387.95 | 93.3 | 11 | 5 | 2.5 | 0.25 |
| 15 | 358.5 | 93.75 | 40 | 5 | 2.5 | 0.25 |
